# Aging remodels the neuronal activity-dependent transcriptome

**DOI:** 10.64898/2026.11.17.767442

**Authors:** Joowon Um, Cecilia Lei, Kodai Nakamura, Eric Tycksen, Jacob W. Adelman, Elizabeth Pollina

**Affiliations:** Dept. of Developmental Biology, Washington University School of Medicine, St. Louis MO, 63110

## Abstract

Experience-driven neuronal activity induces coordinated transcriptional programs in neurons and glia that support plasticity and memory. How aging reshapes these activity-regulated gene (ARG) programs, and whether activity mitigates or exacerbates aging phenotypes, remain open questions with implications for cognitive decline. Here, we develop novel-environment and chemogenetic paradigms to characterize age-dependent remodeling of ARG transcriptomes in the mouse hippocampus at single-cell resolution. Aging contracts youthful ARG programs in both gene number and dynamic range, a contraction that persists under controlled chemogenetic stimulation independent of behavior. In parallel, novel experience in older mice reveals a latent transcriptional competence. First, distinct sets of genes respond to neuronal activity selectively in older cells. Second, enhancing activity in older animals partially reverses, or buffers, basal aging-associated changes across the transcriptome. Novel experience counteracts aging-associated changes in the target genes of the memory regulator CREB1 and the ER-stress sensor ATF6 across cell types. These data constitute a resource for neuronal activity-dependent transcription across the lifespan and identify a basis for the benefits of cognitive engagement: novel experience buffers age-related transcriptional drift.

## Introduction

A fundamental question in aging biology is how cells maintain their adaptive capacity over a lifetime. This challenge is exemplified in the brain, a tissue composed of the body’s longest-lived cells that must continually respond to experience to control learning, memory, and behavior. At the circuit and behavioral level, adaptive capacity declines with age, particularly in the mammalian hippocampus. Hippocampal long-term synaptic potentiation declines in aged rodents^1,2^, and episodic memory and cognitive flexibility are impaired in aged mouse and human brains^3–5^. However, mounting evidence also suggests surprising resilience: aged animals exposed to enriched environments^6–8^ or persistent exercise^9–11^ display partial restoration of decreased synaptic density and cognitive function in memory tasks. In humans, exposure to novelty and cognitive training correlates with improved spatial learning and executive function^12–14^. These studies highlight novel experience as a highly accessible, non-invasive intervention for brain aging. However, the molecular basis of this restored plasticity remains unclear, and how specific conditions can unlock adaptive potential in the aged brain has yet to be fully elucidated.

In the brain, the ability of postmitotic neurons to adapt to new experience is regulated, in part, by the induction of de novo transcriptional programs in response to elevated neuronal activity. Activity-regulated genes (ARGs) play essential roles in synaptic and dendritic growth^15,16^, memory consolidation^17^, adaptive myelination^18^, and genome maintenance^19^. The ARG program is a direct molecular link between experience and lasting changes in neural function. Yet, whether this program remains intact in the aging brain, and whether engaging the ARG transcriptome counteracts aging-associated molecular changes remain unknown. In some studies, canonical immediate-early genes, such as *Fos, Egr1, and Arc,* show reduced mRNA expression with age in rodent models^20–23^ while other studies report no change or even increased *Fos* expression with age^2^. These targeted studies on select genes in rodent models leave open the more fundamental question of how the entire activity-dependent transcriptome changes across the lifespan.

While many multi-omic studies have profiled gene expression under steady-state conditions during aging^24–30^, these approaches cannot reveal the regulatory properties that govern lifetime plasticity in long-lived brain cells, especially those that emerge under stimulation. For example, genome-wide transcriptomic studies conducted under basal conditions in human brains have reported diminished expression of ARGs during aging^24–26^. However, because ARG expression can serve as a readout of neuronal activity levels, these studies cannot distinguish cell types that are simply less active from active cells that are deficient in their ability to induce and regulate ARGs. Furthermore, activity-dependent transcription has been primarily characterized in neurons, despite evidence that astrocytes and other glia mount transcriptional responses to activity^31–35^. To our knowledge, no study has yet examined activity-dependent transcriptional programs across diverse brain cell types over the course of a lifespan. Addressing this gap in knowledge is essential for understanding how experience-based interventions can slow brain aging, and at what point in life they are likely to work.

We therefore designed complementary paradigms to interrogate how the induction and regulation of activity-dependent gene programs change with age: physiological exposure to novel environment (NE) and chemogenetic activation. NE provides a naturalistic stimulus, while chemogenetic (Designer Receptors Exclusively Activated by Designer Drugs (DREADD))^36,37^ activation delivers uniform, time-locked stimulation to neuronal populations, allowing us to resolve age-related changes in ARG inducibility without the variability inherent in behavioral paradigms. We focus on the hippocampus, a region particularly susceptible to aging and neurodegenerative diseases^38^ in which activity is known to drive robust transcriptional responses^19,39–42^. Here, we show that aging remodels, rather than simply attenuates, activity-dependent transcription. Though the classical ARG program induced in young cells is diminished in dynamic range, we also identify an unexpected transcriptional capacity in the aged hippocampus that is revealed by stimulation. Together, these data demonstrate that novel experience reshapes the transcriptome in ways that partially oppose age-dependent alterations.

## Results

### Paradigms to assess activity-dependent gene regulation across aging

We first developed an *in vivo* paradigm to study activity-dependent gene expression in response to physiological stimulation via novel environmental (NE) exposure in mice of different ages (**Fig. 1a**). Conventional NE paradigms, using novel objects and/or running wheels, often activate ARG programs in only sparse cell populations^19,39,43^ and thus limit statistical power to identify transcriptional responses by single cell approaches^19,33,44^. We reasoned that detecting robust differences to ARG transcription during aging required a platform that more uniformly induces ARGs throughout hippocampal cell types and across mice of different ages. We therefore designed an enhanced novel environment paradigm (eNE) inspired by the Marlau cage^45^ that combines features most strongly associated with hippocampal recruitment in a single environment, including a reconfigurable maze, running wheels, tunnels, climbing ramps, and an assortment of novel objects (**Fig. 1a**). We used the expression of canonical ARGs *Fos* and *Npas4* to confirm that exposure to the eNE paradigm drove strong RNA induction in the hippocampus (**Fig. 1b, Extended Data Fig. 1a**). Using this eNE paradigm, we observed markedly stronger and more uniform FOS protein induction in young mice across hippocampal subregions *Cornu Ammonis* (CA) CA1, CA2, CA3 and the dentate gyrus (DG) than a classical NE protocol (**Fig. 1c, Methods**). Widespread FOS expression across brain regions (*e.g.* hippocampus, cortex, thalamus) under the eNE paradigm was independently corroborated by using a *Fos-iCreERT2; Ai14* (Fos-TRAP2) line^46^ (**Extended Data Fig. 1b**). Having validated the eNE paradigm, we collected young, middle-age, and old animals from either home cage (HC) controls or following 6 hours of eNE exposure, a duration that captures both early- and late-response genes, for validation of neuronal activation by immunostaining (left hemisphere) and single-cell RNA-sequencing (right hemisphere) (**Fig. 1a**). We performed these assays on both male and female cohorts to capture the aging biology that can influence both sexes. We confirmed that eNE exposure induced significant FOS protein relative to HC controls in males and females at every age (**Fig. 1d, Extended Data Fig. 1c**). Thus, eNE robustly activates classical activity-dependent gene markers across the lifespan and therefore provides a physiological platform to interrogate age-dependent transcriptional plasticity.

**Figure 1:**
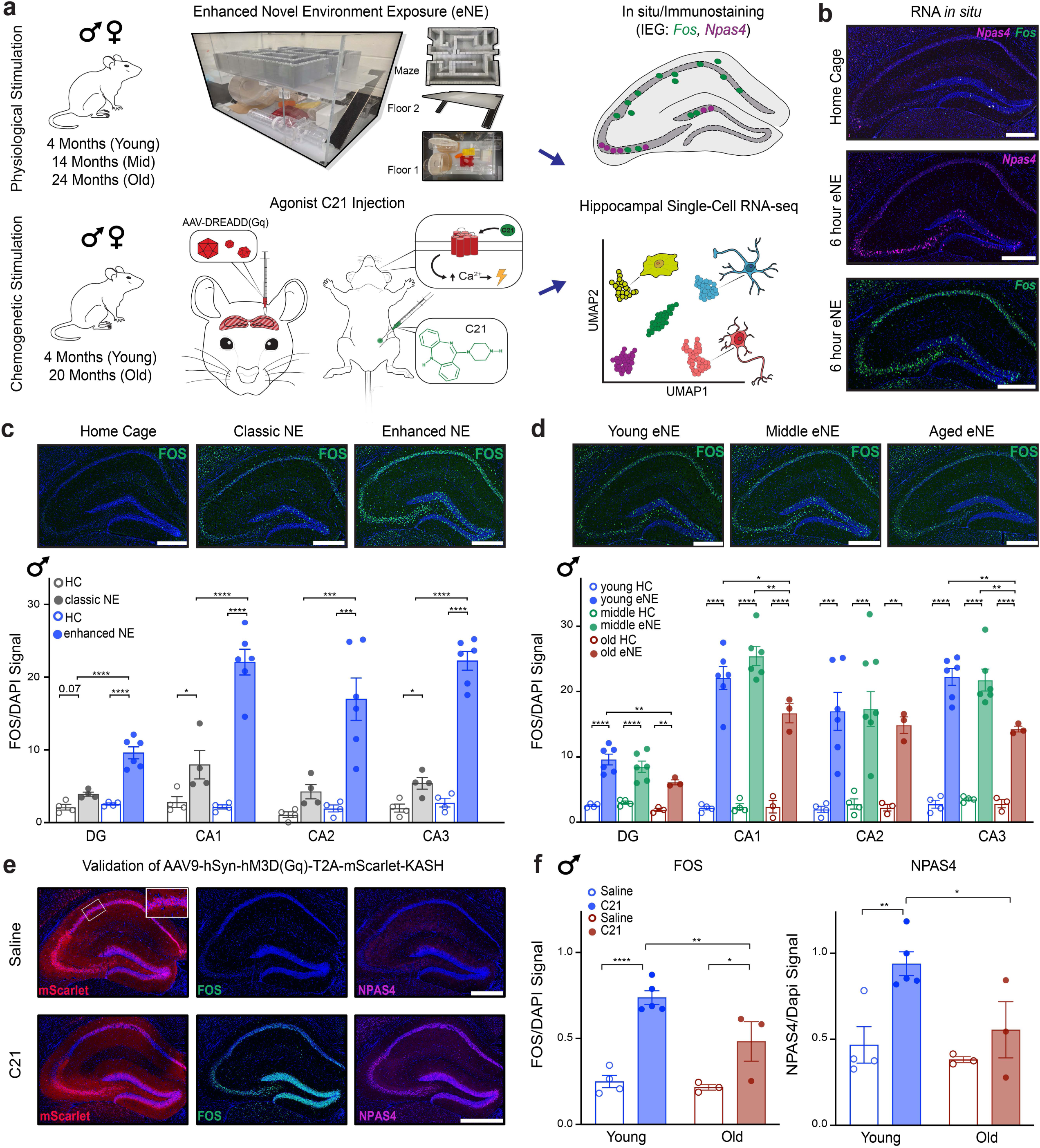
Paradigms to assess activity-dependent gene regulation across aging. a. Schematic of enhanced novel environment (eNE) and chemogenetic activation (DREADD) paradigms to assess neuronal activity in young, middle-age, and old males and females. Samples were collected after 6 hours of eNE exposure or 2 hours of C21 exposure and processed for immunostaining (left hemisphere) or single-cell RNA-seq (right hemisphere). b. *In situ* hybridization for *Fos* and *Npas4* mRNA in home cage control (upper panel) or following 6 hour of eNE exposure. Scale bar indicates 500 µm. c. Upper: Immunostaining for FOS protein in home cage (HC) controls, classical NE paradigm, or newly developed enhanced NE (eNE). Scale bar indicates 500 µm. Lower: Mean ± SEM intensity of FOS immunostaining signal. Each dot represents a single animal as a biological replicate (n=4-6 replicates per group). *p<0.05, **p<0.01, *** <0.005, **** p<0.0001; p-values by 2-way ANOVA (age × stimulation) with Tukey’s multiple hypothesis testing performed within each subregion (*i.e.* DG, CA1, CA2, CA3). d. Upper: Immunostaining for FOS protein in young (4-month), middle age (14-month), or old (24 month) male mice following 6 hours of eNE exposure. Scale bar indicates 500 µm. Lower: Mean ± SEM intensity of FOS immunostaining signal. Each dot represents a single animal as a biological replicate (n=3-6 replicates per group). *p<0.05, **p<0.01,*** p<0.005, **** p<0.0001; p-values by 2-way ANOVA (age × stimulation) with Tukey’s multiple hypothesis testing performed within each hippocampal subregion. e. Immunostaining of NPAS4 and FOS in young male mice injected with AAV9-hSyn-hM3D(Gq)-T2A-mScarlet-KASH. Animals were collected 2 hours post IP injection of DREADD agonist C21 or saline controls. Inset shows nuclear envelope localization of mScarlet-KASH to mark DREADD-expressing nuclei. Scale bar indicates 500 µm. f. Mean ± SEM intensity of FOS immunostaining signal (left) or NPAS4 (right) signal in male mice across the entire hippocampus. Each dot represents a single animal as a biological replicate (n=3-5 replicates per condition). *p<0.05,** p<0.01, **** p<0.001; p-values by 2-way ANOVA (age × stimulation) with Tukey’s multiple hypothesis correction.

In behavioral paradigms that induce ARGs, it remains challenging to fully control the extent to which young and old animals engage with a task or learning paradigm. Thus, as a complement to our eNE paradigm, we established a chemogenetic platform to provide uniform and time-locked stimulation to young and old cells. We designed an AAV viral vector that expresses the excitatory DREADD^36,37^ receptor coupled to a nuclear membrane tagged mScarlet under the neuronal human synapsin (hSyn) promoter (**Fig. 1e**). We delivered DREADD viral constructs into the whole hippocampus of young and old mice. Because differences in DREADD agonist Compound C21 (C21) exposure or viral transduction could confound age comparisons, we verified that C21 levels in the brain, viral spread, and reporter expression levels were equivalent in young and old mice (**Extended Data Fig. 1d-f; Methods**). C21 robustly induced ARGs FOS and NPAS4 in infected cells (**Fig. 1e,f, Extended Data Fig. 1g,h**), with a subtle decline in expression levels in aged brains (**Fig. 1f, Extended Data Fig. 1h**). Together, these platforms provide a conceptual framework to study activity-dependent gene regulation in the aging brain: eNE provides a physiological stimulus relevant to lived experience, while DREADD activation delivers time-locked input that isolates age-related changes in ARG induction from differences in behavioral engagement.

### Aging remodels the activity-dependent transcriptome with uniquely vulnerable cell types

Focusing first on our physiological eNE paradigm, we sought to examine whether the activity-dependent program is altered with age and in which types of cells age has the most significant effects. We generated a single-cell atlas of the aging hippocampus from 36 mice: young, middle-aged, and old animals of both sexes under basal (home cage, HC) conditions or following 6 hours of eNE, with 3 biological replicates per condition. This yielded 818,098 high-quality cells spanning 18 cell types (**Fig. 2a, Extended Data Fig. 2**), which we annotated manually using canonical neuronal and glial markers (**Extended Data Fig. 3**). We employed the 10X Genomics Chromium Flex assay, a multiplex probe-based method of RNA profiling from fixed samples. Multiplexing allows ages and conditions to be processed together to minimize batch effects, while sample fixation preserves the transcriptional state at collection and thus reduces dissociation-induced artifacts. Consistent with this, cells co-embedded across ages, sexes, and stimulation conditions (**Fig. 2a**), indicating minimal batch effects.

**Figure 2:**
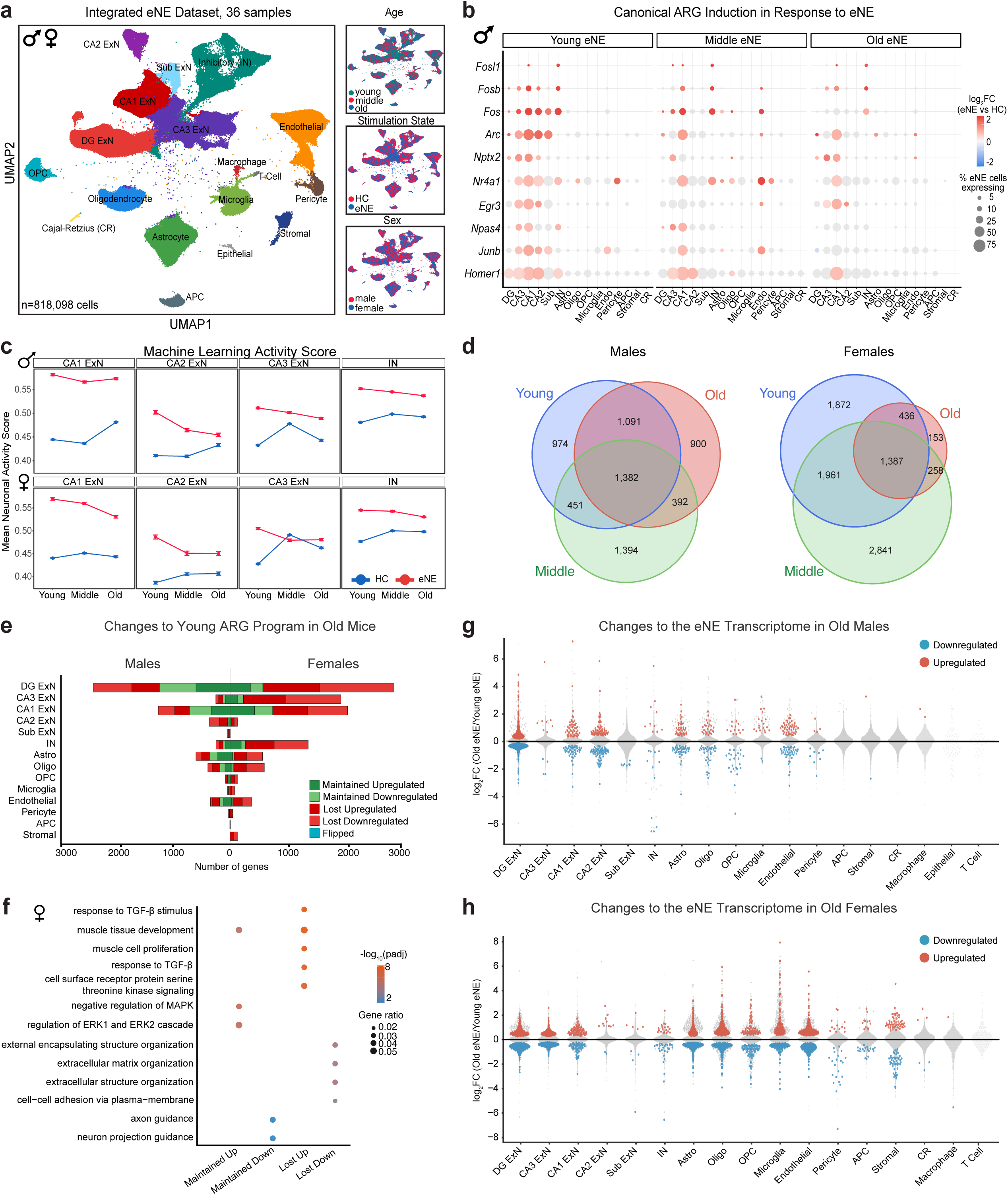
Aging remodels the ARG transcriptome with uniquely vulnerable cell types. a. UMAP of 818,098 hippocampal cells from 36 young, middle-aged, and old male and female mice housed in the home cage (HC) or exposed to an enhanced novel environment (eNE) for 6 h (n = 3 mice per sex, age, stimulation state). Insets show cells colored by age, stimulation state, and sex. Cell types were annotated using canonical markers (**Extended Data Fig. 3; Methods**). b. Dot plot of canonical ARGs across cell types in young, middle-aged, and old male mice. Dot color indicates log_2_ fold change (eNE vs HC), and dot size indicates the percentage of eNE cells expressing the gene. Grey dots indicate non-significant changes (padj ≥ 0.05). Females are shown in **Extended Data Fig. 4a**. IN, inhibitory neuron; Sub, subiculum; Astro, astrocyte; oligo, oligodendrocyte; OPC, oligodendrocyte precursor cell; Endo, endothelial cell; APC, astrocyte progenitor cell; CR, Cajal–Retzius cell. c. Mean NEUROeSTIMator activity score in CA1, CA2, and CA3 excitatory neurons and inhibitory neurons from HC (blue) and eNE (red) mice across ages, in males (top) and females (bottom). Error bars indicate SEM across pseudobulk samples. d. Venn diagrams showing overlap of ARGs between young, middle-aged, and old males (left) and females (right). ARGs were defined as genes significantly up- or downregulated by eNE relative to age-matched HC (padj < 0.05, no fold-change threshold) and were pooled across cell types. e. Fate of young ARGs in old mice per cell type (males left, females right). Young ARGs were classified by their regulation in old eNE vs. old HC mice: Maintained Upregulated and Maintained Downregulated (significant in the same direction in old mice), Lost Upregulated and Lost Downregulated (no longer significant), and Flipped (significant in the opposite direction in young vs. old). f. Gene Ontology (Biological Process) enrichment of Maintained and Lost ARG classes from **e** in females, pooled across cell types. Dot color indicates −log_10_(padj), and dot size indicates gene ratio. Males are shown in **Extended Data Fig. 4f**. g,h. Strip volcano plots of differential expression between old eNE and young eNE mice per cell type in males (**g**) and females (**h**). Genes significantly different between old HC and young HC mice were excluded, isolating differences specific to the activated state. Red and blue indicate genes significantly higher or lower, respectively, in old eNE relative to young eNE mice (padj < 0.05); grey indicates non-significant genes. ExN, excitatory neuron; IN, inhibitory neuron; Sub, subiculum; Astro, astrocyte; oligo, oligodendrocyte; OPC, oligodendrocyte precursor cell; Endo, endothelial cell; APC, astrocyte progenitor cell; CR, Cajal–Retzius cell.

With this resource, we first asked whether our eNE paradigm drove an activity-dependent response capturing canonical ARGs across hippocampal cells throughout lifespan (**Fig. 2b, Extended Data Fig. 4a**). We employed a pseudobulk approach using DeSEQ2^47^ to quantify differential gene expression between age-matched HC controls and eNE samples, using each animal as an independent replicate^48,49^. Because rare populations yielded low cell counts in some samples, cell types with fewer than two pseudobulk samples were excluded; as a result, not all cell types are represented in both sexes (**Methods**). In young mice, experience-driven neuronal activity triggered transcription in neuronal and glial compartments throughout the hippocampus, with a substantial induction in the principal excitatory populations, DG and CA1-3 pyramidal neurons (**Fig. 2b, Extended Data Fig. 4a,b**). In line with published studies, the young-induced program in neurons (*e.g.* CA1) was enriched for synaptic plasticity, kinase, and growth factor signaling^34^ while ARGs in astrocytes were enriched for metabolic changes and response to glucocorticoid^50^ (**Extended Data Fig. 4c,d**). Thus, eNE in young animals drives a functionally coherent transcriptional program that serves as a reference against which to measure age-related change.

We examined age-dependent changes to known immediate early gene transcription factors (*e.g. Fos, Npas4*), which coordinate downstream transcriptional responses (**Fig. 2b**). We observed that during aging, the number and degree of induction for these canonical ARGs was reduced in most cell types (**Fig. 2b, Extended Data Fig. 4a**). To corroborate these findings with an independent analysis, we applied NEUROeSTIMator, a machine learning model trained on the expression of canonical ARGs to generate estimates of activity state in individual cells^51^. Single-cell activity scores demonstrate activation in eNE across neuronal cell types and reveal a consistent decrease in old animals (**Fig. 2c**).

We next quantified the breadth of the inducible program as the number of genes significantly regulated by eNE relative to age-matched HC across all cell types in young, middle-aged, and old animals (**Fig. 2d**). While a core set of ARGs was reproducibly induced at all ages, the total number of ARGs was reduced in old animals. We then decomposed the youthful ARG program into genes maintained or lost with age across all hippocampal cell types (**Fig. 2e**). This contraction was severe in females. For example, old females lost 83% of youthful ARGs in CA3, 80% in dentate gyrus, 87% in astrocytes, and 89% in oligodendrocytes. In males, the contraction was more restricted, with roughly half of youthful ARGs lost across most cell types and the largest losses in CA2 (89% no longer induced). Notably, middle-aged females retained more youthful ARGs than old females in every cell type examined, suggesting that late-life changes drive the deficit in activity-dependent responses in females (**Extended Data Fig. 4e**).

By gene ontology analysis, maintained genes in both sexes were enriched for the regulation of kinase activity, likely reflecting that the upstream signaling that drives initial ARG induction tends to be preserved with age (**Fig. 2f, Extended Data Fig. 4f**). Among upregulated ARGs, the genes no longer induced in old females were enriched for the cellular response to TGF-β. In old males, the cellular response to TGF-β was preserved while genes no longer induced are especially enriched in protein folding and processing. Across sexes, downregulated ARGs lost with age were associated with ECM organization and deposition, processes known to stabilize synapses and restrict remodeling^52,53^; thus, the transcriptional downregulation of ECM modifying genes that contributes to activity-induced structural plasticity in youth is attenuated in aged animals (**Extended Data Fig. 4f**).

To identify differences in activated states between young and old cells, we directly compared old eNE to young eNE (**Fig. 2g,h**). We specifically excluded genes that significantly differed under basal conditions to focus on how activity can reveal changes in aged cells (**Fig. 2g,h, Methods**). In males, age-dependent changes in old eNE relative to young eNE were concentrated in the DG, with few changes in any other population. In females, the changes were more broadly distributed: CA3 pyramidal neurons exhibited the most changes, in addition to astrocytes and oligodendrocytes. Even at middle-age, we observe significant differences in the eNE transcriptome relative to young, with DG especially affected in males (**Extended Data Fig. 4g,h**). At middle age, non-neuronal cells showed the highest numbers of DEGs, indicating that glia are among the first populations to shift their activated state during aging (**Extended Data Fig. 4g,h**).

We next sought to gain improved cellular resolution for populations of cells known to be highly heterogenous in their responses to activity, including both inhibitory neurons (INs) and astrocytes. Subclustering resolved four distinct IN populations – somatostatin (SST), parvalbumin (PV), vasoactive intestinal peptide (VIP) and lysosomal associated membrane protein family member 5 (LAMP5) – each of which showed distinct activity-dependent responses across age and sex (**Extended Data Fig. 5a-c**). IN subclusters induced canonical ARGs (**Extended Data Fig. 5d**) as well as a program of inhibitory-enriched ARGs not present in excitatory neuron populations (**Extended Data Fig. 5e**). Among IN subtypes, PV neurons display the greatest age-related loss in females, as evidenced by the relatively higher number of DEGs when comparing young vs. old eNE states (**Extended Data Fig. 5f**) and a substantial loss of the youthful ARG repertoire (**Extended Data Fig. 5g**). By performing a similar subcluster analysis among astrocytes (**Extended Data Fig. 6a**), we identified distinct ARG profiles across the subclusters, with the predominant C1 cluster as the most transcriptionally responsive to eNE (**Extended Data Fig. 6b-g**). Together, these analyses define new ARG genes across both neurons and glia and suggest vulnerability in the ARG-program during aging is highly cell-type specific.

### Reduced dynamic range in youthful ARGs is a main mode of failure with age

We hypothesized that the contraction of the youthful ARG program across lifespan is due, in part, to a reduction in the dynamic range of gene inducibility, thereby reducing the ability of the cell to mount a graded response to stimuli. To visualize these age-related dynamics, we tracked the log_2_ fold change (FC) of youthful ARGs in middle-age and old cohorts (**Fig. 3a,b, Extended Data Fig. 7a,b**). We observed a progressive attenuation in the magnitude of the transcriptional response, with genes exhibiting significantly higher log_2_FC values in young animals compared to both middle-aged and old cohorts. This trend was consistent across both up-regulated and down-regulated gene sets, suggesting that aging imposes a bidirectional dynamic range compression on the hippocampal activity-dependent program. Importantly, this dampening effect was not uniform throughout lifespan; a reduction in fold change commonly occurred between young and middle-age, identifying mid-life as an inflection point for the attenuation of neuronal and glial activity-dependent responses. Notably, we identified microglia in females as the cell type with the most dynamic range compression (both up and downregulated genes) (**Extended Data Fig. 7c**).

**Figure 3:**
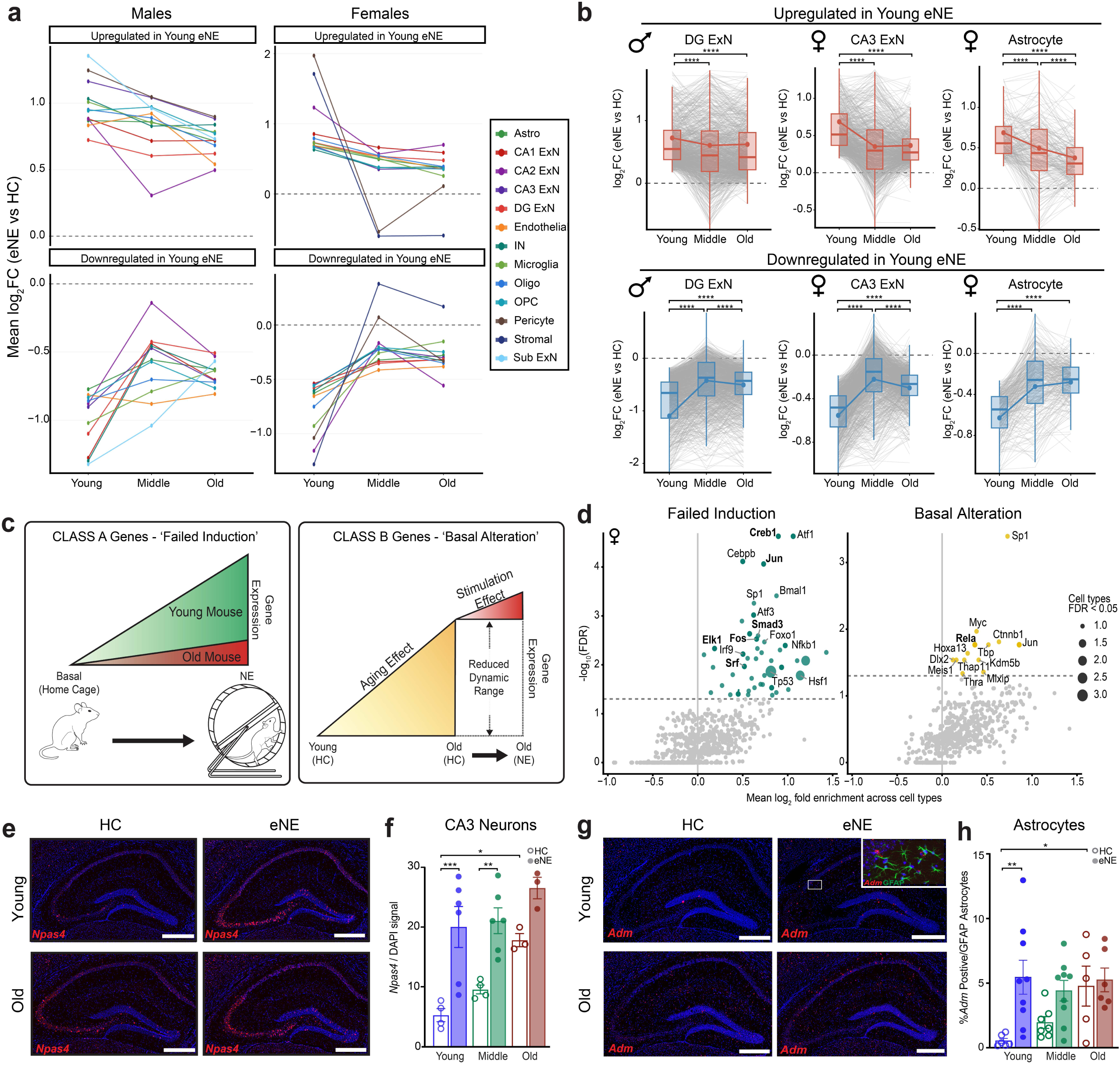
Reduced dynamic range in youthful ARGs is a main mode of failure with age. a. Mean log_2_FC (eNE vs. HC) of young ARGs in young, middle-aged, and old mice in males (left) and females (right). Young ARGs were genes significantly upregulated (top) or downregulated (bottom) in young eNE vs. young HC mice (padj < 0.05, no fold-change threshold). Young ARG log_2_FC was tracked in the middle-age and old comparisons. Lines connect mean values across ages. Only cell types with ≥ 10 young ARGs in a given direction are shown. b. Per-gene log_2_FC (eNE vs. HC) of young-upregulated (top) and young-downregulated (bottom) ARGs across age in male DG ExNs, female CA3 ExNs, and female astrocytes. Boxplots show the median and interquartile range, with whiskers extending to 1.5× IQR and outliers not displayed. Grey lines connect each gene’s log_2_FC across ages; the y-axis is set to the boxplot range, so only genes falling within that range are drawn. \*\*\*\**p* < 0.0001; two-sided, paired Wilcoxon signed-rank test on all genes in the set, Benjamini–Hochberg corrected for multiple age comparisons within each cell type. c. Schematic of the two modes of ARG failure with age. Class A, failed induction: eNE fails to induce the gene to its young level, without a comparable change in basal expression. Class B, basal alteration: aging shifts basal (HC) expression in the direction of the young eNE response, by at least half its magnitude, so the remaining dynamic range is reduced. d. CollecTRI transcription factor regulons enriched among failed-induction (left) and basal-alteration (right) genes in females. The x-axis shows the log_2_ fold enrichment of each regulon averaged across cell types; the y-axis shows −log_10_(FDR). Dot size indicates the number of cell types in which the TF was significant (FDR < 0.05). The dashed line marks FDR = 0.05. Males are shown in **Extended Data Fig. 7e**. e. Representative RNAscope in situ hybridization for *Npas4* (red) with DAPI (blue) in the hippocampus of young and old HC vs. eNE mice. Scale bars, 500 µm. f. *Npas4* signal normalized to DAPI in CA3 neurons of young, middle-aged, and old HC (open) and eNE (filled) male mice. Females are shown in **Extended Data Fig. 7f**. g. Representative RNAscope for *Adm* (red) with DAPI (blue) in young and old HC vs. eNE mice. The inset shows *Adm* with GFAP immunostaining (green) in the boxed region. Scale bars, 500 µm. h. Percentage of GFAP⁺ astrocytes positive for *Adm* in young, middle-aged, and old HC (open) and eNE (filled) male mice. Females are shown in **Extended Data Fig. 7g**. For f and h, data are mean ± SEM, and each dot represents one mouse. \**p* < 0.05, \*\**p* < 0.01, \*\*\**p* < 0.001; two-way ANOVA (age × stimulation) with Tukey’s multiple comparisons test.

To understand the mechanisms driving this compression, we examined two routes of transcriptional failure: (1) failed induction, a blunting of the stimulus-evoked response, and (2) basal age effects, such as a de-repression of the gene under unstimulated conditions that reduces the available range for activity-induced expression change (**Fig. 3c, Methods**). We note that most compromised genes are likely affected by a combination of both. We quantified the relative contribution of the two mechanisms across cell types. Failed induction was the pre-dominant route of failure, though basally altered genes were identified across most cell types in both sexes (**Extended Data Fig. 7d**). To identify the regulatory basis of failed-induction and basal alteration, we performed transcription factor activity inference (CollectTRI)^54^ on each gene category (**Fig. 3d, Extended Data Fig. 7e**). As expected, failed induction genes were highly enriched for targets of known activity-modified transcription factors such as CREB1, SRF, EGR1 (**Fig. 3d, Extended Data Fig. 7e**), as well as SMAD3, consistent with the loss of TGF-β signaling in old females (**Fig. 2f**). However, in both sexes, basal alteration genes were enriched for inflammatory signaling transcription factors such NFkB, RELA, and STAT1 (**Fig. 3d, Extended Data Fig. 7e**). This finding suggests that age-dependent pro-inflammatory signals drive essential changes to the basal aging transcriptome that thereby alters the activity-dependent response^55^. This result adds to our growing understanding of the detrimental effects of ‘inflammaging’^55^, which may disrupt youthful plasticity mechanisms by interfering with activity-dependent transcription. To validate the phenomenon of basal alteration by an orthogonal method, we performed *in situ* hybridization for the canonical ARG *Npas4* (**Fig. 3e,f, Extended Data Fig. 7f**) and the novel astrocyte-inducible gene *Adm*, both of which showed elevated resting expression in aged tissue (**Fig. 3g,h, Extended Data Fig. 7g**). Together, these data show that aging does not simply switch off activity-dependent transcription but compresses its dynamic range through two routes, failed induction and basal alteration, which are deployed differentially across genes and vulnerable cell types.

### Dynamic range compression is independent of behavioral engagement

Our data has thus far revealed a progressive contraction of the youthful canonical ARG program. However, novel environment cannot equalize neuronal activity across ages, leaving unresolved whether these changes reflect differences in behavior at the organismal level and/or downstream consequence of reduced upstream drive in aged circuits. Chemogenetic activation removes these confounds, imposing a time-locked stimulus directly on young and old cells (see **Fig.1**). We focused our analysis on females, in which the enrichment response was notably impaired with age (**Figs. 2-3**). We profiled the hippocampus from agonist C21 versus saline-treated young and old female mice using the same 10X Genomics FLEX assay. The resulting dataset, comprised of 3 samples per condition, co-embedded into a well-integrated manifold with 17 unique neuronal and non-neuronal clusters (**Fig. 4a,b**). Our design incorporated a custom probe set targeting the mScarlet reporter, enabling DREADD-infected cells to be identified directly from the transcriptome (**Fig. 4a,b**). Reporter-expressing cells were distributed across the neuronal clusters (**Fig. 4b**, right), consistent with hSyn-driven, neuron-enriched DREADD expression.

**Figure 4:**
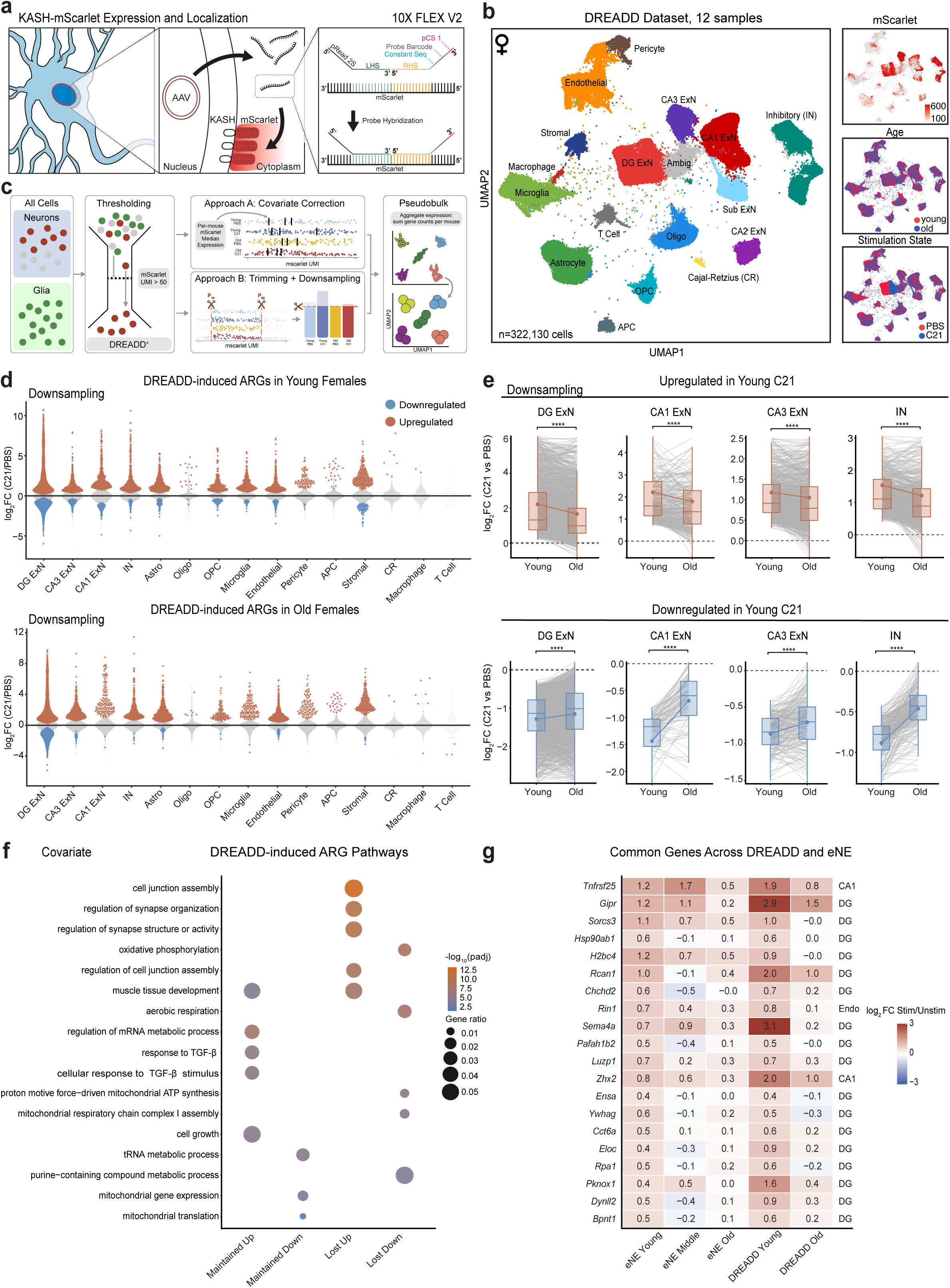
Dynamic range compression is independent of behavioral engagement. a. Schematic of the DREADD construct and detection strategy. AAV9-hSyn-hM3D(Gq)-T2A-mScarlet-KASH drives neuronal expression of the hM3Dq receptor and an mScarlet reporter anchored to the outer nuclear membrane by a KASH domain, marking the nuclei of DREADD-expressing neurons. Custom 10x Flex probe pairs targeting mScarlet transcripts allowed transgene detection in single cells. b. UMAP of 322,130 hippocampal cells from young and old female mice treated with C21 or PBS (n = 3 mice per age × stimulation). Right: mScarlet UMI counts (cells with ≥ 50 UMIs), age, and stimulation state overlaid on the UMAP. c. Schematic of the analysis pipeline. Neurons expressing mScarlet were identified, and two strategies, covariate correction and downsampling, were used to match viral transduction across mice before pseudobulk analysis (see **Methods**). d. Strip plots of differential expression between C21 and PBS in young (top) and old (bottom) females. Red and blue indicate significantly up- and downregulated genes (padj < 0.05); grey indicates non-significant genes. ExN, excitatory neuron; IN, inhibitory neuron; Sub, subiculum; Astro, astrocyte; Oligo, oligodendrocyte; OPC, oligodendrocyte precursor cell; Endo, endothelial cell; APC, astrocyte progenitor cell; CR, Cajal–Retzius cell. e. Per-gene log_2_FC (C21 vs. PBS) of young-upregulated (top) and young-downregulated (bottom) ARGs in female DG ExNs, CA1 ExNs, CA3 ExNs and INs. Boxplots show the median and interquartile range, with whiskers extending to 1.5× IQR and outliers not displayed. Grey lines connect each gene’s log_2_FC across ages; the y-axis is set to the boxplot range, so only genes falling within that range are drawn. \*\*\*\**p* < 0.0001; two-sided, paired Wilcoxon signed-rank test on all genes in the set, Benjamini–Hochberg corrected for multiple age comparisons within each cell type. f. Gene Ontology (Biological Process) enrichment of DREADD-induced ARG classes: maintained and lost, up- and downregulated genes. Dot color indicates −log_10_(padj) and dot size indicates gene ratio; the top five terms per class are shown. g. Genes induced by both eNE and DREADD in young mice whose response is compressed with age in both paradigms. Values are log₂FC (stimulated vs. unstimulated) at each age, averaged across the cell type in which the gene was compressed; those cell types are listed on the right. The 20 most strongly compressed genes are shown. Unless otherwise noted, DREADD analyses used the downsampling approach; results with the covariate approach are shown in **Extended Data Fig. 8**.

Importantly, the samples selected for single-cell RNA-seq displayed no significant differences in mScarlet intensity or spread based on tissue-level immunostaining, and we detected equivalent levels of C21 in the cortex of young and old mice (**Extended Data Fig. 8a,b**). Because the recovery of mScarlet-expressing cells could differ across samples and bias differential-expression counts, we restricted analysis to cells with bona fide reporter expression (50 raw mScarlet UMI per cell). We further normalized for mScarlet levels using two independent strategies, a per-mouse covariate and downsampling to equalize mScarlet+ cells per cell type (**Fig. 4c, Extended Data Fig. 8c-e, Methods**). Post-filtering, mScarlet distributions were matched across the cohort (**Extended Data Fig. 8d,e**), indicating that any age difference in induction reflects the neuronal response rather than differential stimulation or capture.

Relative to saline-controls, C21 induced a broad transcriptional program across neuronal and non-neuronal cell types in young neurons and, critically, in aged neurons (**Fig. 4d**). Thus, aged neurons remain competent to mount an activity-dependent response to direct, uniform stimulation. Though we did observe many gene expression changes in non-neuronal cells in both young and old mice, we focus hereafter on the neuronal responses during aging, as we cannot as precisely control stimulation levels in the glia. Although aged neurons mounted a diverse transcriptional response to C21, its composition differed substantially from young neurons, with loss of the youthful program concentrated in DG, CA3, and INs (**Extended Data Fig. 8f**). Using both our co-variate and downsampling approaches, we found that log_2_FC distributions comparing saline to C21 for youthful ARGs displayed bidirectional compression with age in most neuronal populations (**Fig. 4e, Extended Data Fig. 8g**), a result that mirrors the dynamic range compression observed under eNE (**Fig. 3a,b**). Pathway analysis of genes no longer significantly induced by DREADDs in old females especially highlight a failure of synaptic plasticity associated genes (**Fig. 4f**), a subset of which similarly lose their inducibility in eNE (**Fig. 4g**). We next ranked genes by the extent of their age-related loss in inducibility across both DREADD and eNE paradigms. The most compressed genes were not canonical immediate early genes but second-wave effectors, particularly enriched in DG ExNs, that regulate synaptic plasticity (*Sorcs3*, *Rin1*, *Rcan1*, *Gipr*), neuronal transport (*Dynll2*, *Pafah1b2*, *Ywhag*), protein folding (*Hsp90ab1*, *Cct6a*), and genome maintenance (*Rpa1*) (**Fig. 4g**). Together, these results indicate that aging attenuates the full dynamic range of the core ARG response under direct chemogenetic stimulation and demonstrate that the contraction of the activity-dependent transcriptional program with age is, in part, independent of behavior.

### Aging cells retain a latent capacity to remodel the transcriptome in response to activity

Our data thus far suggest that aging contracts the youthful activity-dependent transcriptional program in dynamic range, beginning even at middle age (**Fig. 2-3**). However, we observed that both middle-aged and old animals regulate a set of ARGs not detected in young animals (**Fig. 2d, Extended Data Fig. 9a**), suggesting that older cells do retain a latent competence to uniquely respond to novel environment. In the female cohort, age-emergent ARGs were broadly distributed across cell types at middle age but far more restricted in number and cell type distribution in old animals (**Extended Data Fig. 9a**). Age-emergent genes in old animals were highly enriched for synapse assembly and immune signaling (**Extended Data Fig. 9b,c,d**) and included targets of the known longevity-associated factor FOXO1^56^, the cholesterol synthesis regulator SREBF2^57^, and the anti-inflammatory hormone receptor PGR^58^ as well as classical ARG transcription factors (**Extended Data Fig. 9e**). Activity in aged cells thus engages regulators of metabolic and neuroprotective pathways that could represent a mechanism to compensate for age-related changes in metabolism and inflammatory state^27,55^.

The observation that in response to novel environment, aged cells regulate a set of genes that counteract common aging phenotypes (for example, upregulating synaptic genes that decline with age in neurons^27^ while downregulating age-induced inflammatory pathways in glia^27,55^) prompted a broader question: does novel experience shift the aged transcriptome back toward the youthful state? Thus, we asked whether experience-induced neuronal activity can buffer the broader transcriptional drift associated with aging (**Fig. 5a**). For each cell type, we compared the basal aging transcriptional vector (old HC vs. young HC) with the activity vector at the same age (old eNE vs. old HC) by cosine similarity; negative values denote directional opposition, and positive values denote alignment with aging. We ensured that the aging and activity vectors, both of which use the HC controls as a reference, were not directly coupled. We therefore calculated the cosine using an iterative ‘held-out’ approach such that the aging and activity vectors were estimated from independent HC animals (**Extended Data Fig. 10a**). We further validated the cosine measurement using an independent approach, which accounts for mouse-to-mouse variation in HC animals (**see Methods for both approaches**). At middle age, eNE opposed the aging vector in most cell types of both sexes (**Fig. 5b, Extended Data Fig. 10b**). Interestingly, this buffering capacity declined with age, with a marked deficit in females. Indeed, cosine similarly converged toward zero between middle and old age in nearly all cell types in females, with CA1 ExNs as the exception (**Fig. 5b, Extended Data Fig. 10c**). To illustrate what buffering looks like at the gene level, we plotted the aging and activity log₂FC for buffered genes (**Fig. 5c, Methods**). Across cell types, these profiles formed near mirror images; eNE repressed genes elevated by aging and induced genes repressed by aging, as seen by example heatmaps shown for DG, CA3, microglia, and astrocytes (**Fig. 5c, Extended Data Fig. 10d**). For example, eNE counteracted several hallmark features of brain aging by boosting expression of homeostatic microglia markers (*Selplg*, *Nrros*, *Mfge8*) while repressing inflammatory mediators (*Map3k8*, *Fgl2*) in microglia. Likewise, novel experience suppressed the age-associated interferon response in astrocytes (*Oasl2*, *Bst2*, *Ddx60*) and boosted synaptic transmission genes in CA3 ExNs (*Syt2*, *Slc17a6*, *Nts*) (**Extended Data Fig. 10d**). Thus, novel experience can directionally oppose basal aging changes across cell types, though this capacity is compromised in old age.

**Figure 5:**
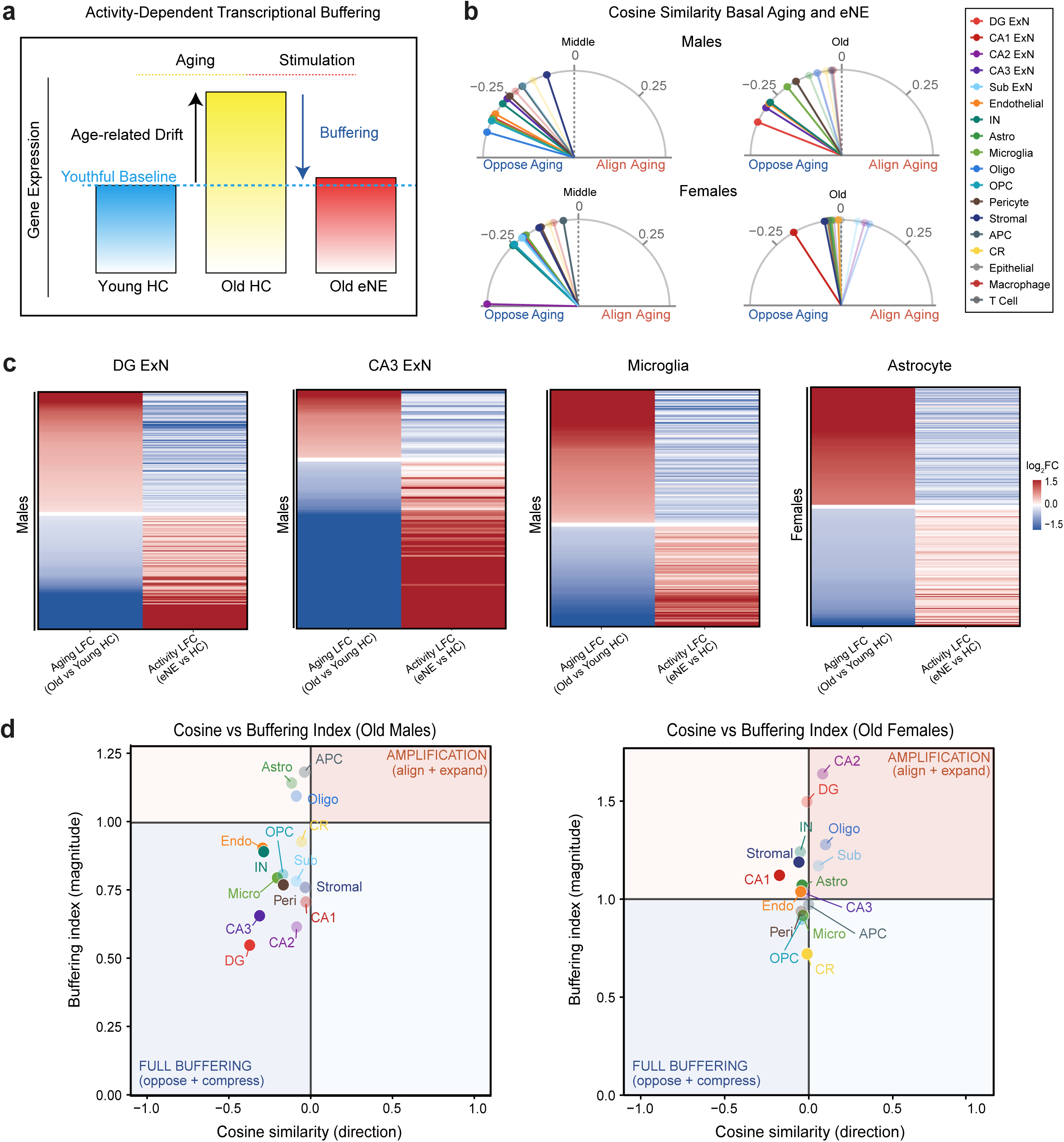
Aging cells retain a latent capacity to remodel the transcriptome in response to activity. a. Schematic of activity-dependent transcriptional buffering. Aging shifts basal gene expression away from the youthful baseline (age-related drift, young HC to old HC). Buffering is the extent to which eNE in aged animals returns expression toward that baseline. b. Cosine similarity between the aging change (aged HC vs. young HC) and the eNE response (eNE vs. HC at the same age) per cell type, at middle (left) and old (right) age in males (top) and females (bottom). Each line is one cell type, with its angle scaled to the cosine value. Lines left of the dashed line (cosine < 0) indicate that eNE opposes aging. Dark symbols mark cell types whose buffering was supported by two independent estimates of the cosine (held-out and bias-corrected); faded symbols mark those supported by only one (**Methods**). c. Buffered genes in male DG ExNs, CA3 ExNs and microglia and female astrocytes. For each gene, the left column shows the aging change (old HC vs young HC), and the right column shows the eNE response in old mice (old eNE vs old HC) (see **Methods, Tier 2 genes**). Genes are split by aging direction and ordered by aging log_2_FC. d. Held-out cosine similarity versus buffering index (BI) per cell type at old age in males (left) and females (right). BI = Σ log2FC²(old eNE vs young HC) / Σ log2FC²(old HC vs young HC). BI < 1 indicates that old eNE mice are closer to young HC mice than old HC mice are. In the lower-left quadrant (cosine < 0, BI < 1, full buffering), eNE both opposes aging and moves expression toward the young state. In the upper-right quadrant (cosine > 0, BI > 1), eNE amplifies aging changes. In the upper-left quadrant (cosine < 0, BI > 1), eNE opposes aging but overshoots the young baseline. Dark symbols mark cell types whose buffering was supported by two independent estimates of the cosine (held-out and bias-corrected); faded symbols mark those supported by only one (**Methods**).

How close does novel experience bring the transcriptome back towards a youthful baseline level? To quantify the extent to which eNE exposure returned the aged transcriptome toward youth, we calculated a buffering index, defined as the distance between old eNE and young HC expression, relative to the distance between old HC and young HC expression (Buffering index = Σ(O_NE − Y_HC)² / Σ(O_HC − Y_HC)²). Values below 1 indicate that eNE exposure brought the aged transcriptome closer to youth whereas values above 1 indicate further away (**Extended Data Fig. 10e**). Cell types with a cosine below 0 and a buffering index below 1 were considered fully buffered: their activity response both opposed aging in the directionality of change and brought the transcriptional levels closer to the youthful baseline state (**Fig. 5d**). In aged males, most cell types met both criteria, with DG ExNs showing the strongest renormalization (**Fig. 5d, Extended Data Fig. 10e**). In aged females, however, most cell types had buffering indices above 1 with cosines near zero, indicating that across the broader transcriptome, novel experience does not oppose aging in females (**Fig. 5d, Extended Data Fig. 10e**). Notably, direct chemogenetic activation of neurons likewise failed to buffer the aging transcriptome in aged females, as most cell types showed cosines near or above zero and buffering indices greater than 1. These results suggest that chemogenetic activation of neurons alone may not be sufficient to drive a transcriptome-wide buffering response, which likely requires the complex signaling that occurs with experience-driven neuronal activity (**Extended Data Fig. 10f**). Although the female transcriptome, as a whole, was not buffered by novel experience in old age, individual genes still displayed buffering, such as the interferon response in aged astrocytes (**Fig. 5c, Extended Data Fig. 10d**). Therefore, experience-induced neuronal activity can still reverse specific age-related programs to restore select aspects of youthful gene regulation. Consistent with the idea that activity can promote youthful signaling during aging, we performed Neuron and Cell Chat^59,60^ analysis to infer the communication strength between different cell types in the hippocampus based on the expression of receptor ligand pairs and found that experience-induced activity partially restores the age-associated decline in signaling potential between hippocampal cell types (**Extended Data Fig. 10g**).

### Distinct transcription factor programs underlie activity-dependent buffering

Finally, we sought to identify potential regulators of activity-dependent buffering, and to determine whether these factors were shared across cell types. Neuronal activity drives transcription through a hierarchy of transcription factors (TFs), many of which are themselves altered with age. We therefore inferred TF activity from the expression of their target genes using a univariate linear model with CollectTRI regulons^54^. We identified TFs whose activity changed with aging under basal conditions and was shifted in the opposite direction by novel experience in old animals (buffered), and TFs for which novel experience pushed activity in the same direction as aging (amplified) (**Fig. 6a**). In old males, the top buffered TFs across cell types were regulators of activity-dependent transcription and cellular stress adaptation, including CREB1 and the ER-stress sensor ATF6, together with a cluster of circadian clock factors (BMAL1, CLOCK) (**Fig. 6a**). These factors were buffered in opposite directions. CREB1 activity declined with age and was restored by eNE, whereas the clock factors gained activity with age and were suppressed by eNE. Stimulation thus restored a lost activity-dependent program while damping an aberrant rhythmic one. In old females, CREB1 was again among the most strongly buffered TFs, together with regulators of mitochondrial biogenesis (NRF1) and proteostasis (HSF1) (**Fig. 6a**). In both sexes, amplified TFs in multiple cell types involved inflammatory and stress-responsive factors, including the glucocorticoid receptor NR3C1 in females, and NF-κB, and steroid hormone receptors in males. These results suggest that novel experience partially restores activity-dependent and homeostatic programs while further engaging stress and inflammatory signaling that aging has already set in motion.

**Figure 6:**
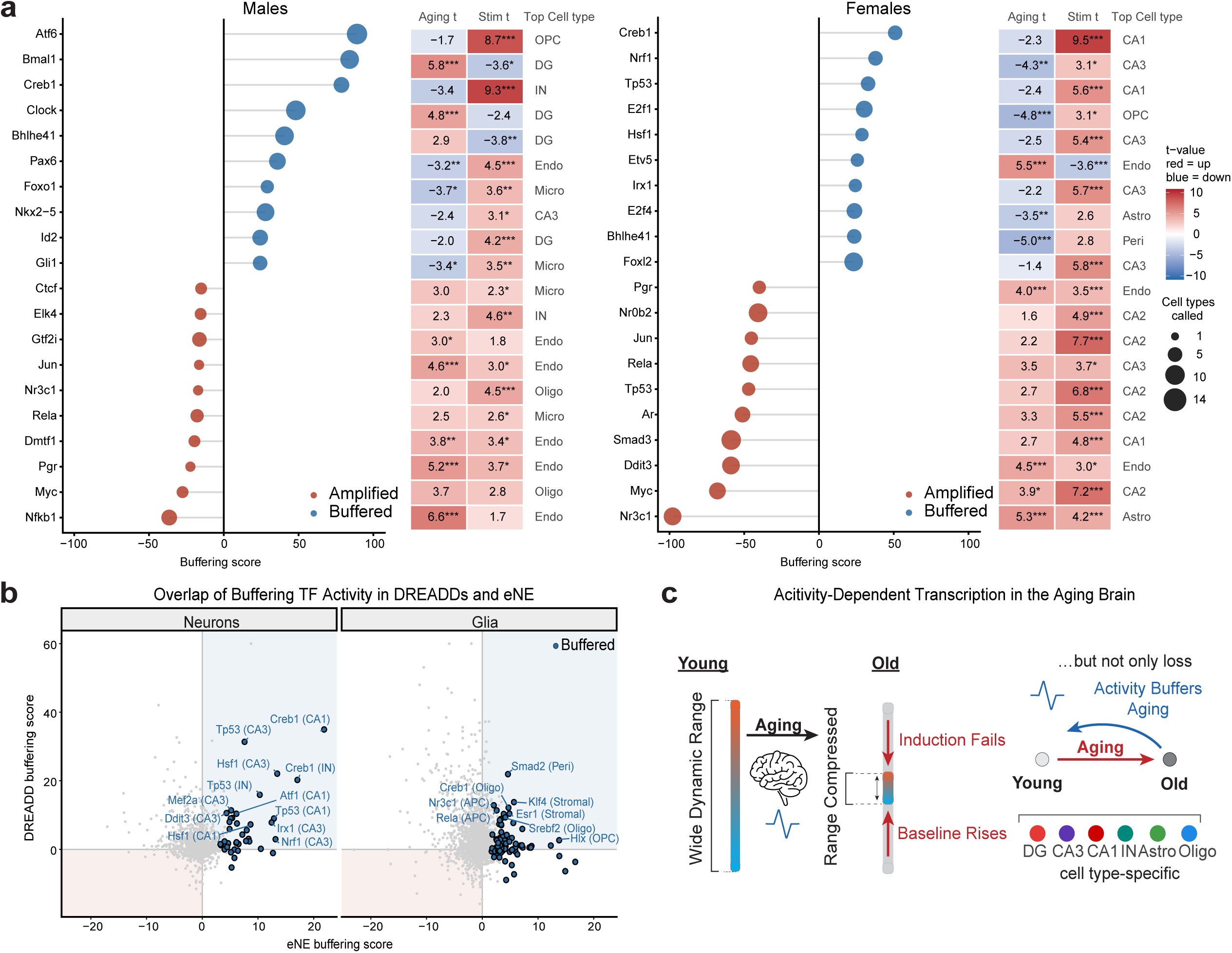
Distinct transcription factor programs underlie activity-dependent buffering. a. Transcription factors (TFs) whose activity is most strongly buffered (blue) or amplified (red) by eNE in old males (left) and old females (right). The buffering score measures how strongly eNE opposes (positive) or reinforces (negative) the aging change in a TF’s activity (**Methods**). TFs are ranked by their score, summed across the cell types in which they were called. Dot size indicates how many cell types contribute to the score. Tiles show the aging and stimulation activity (t-values from ULM) in the cell type with the highest score, named on the right. Stars indicate how confidently each TF activity t-value is estimated: *FDR < 0.05, **FDR < 0.01, ***FDR < 0.001. b. DREADD buffering score plotted against eNE buffering score in old females for all TFs that overlap between DREADDs and eNE. Each point is one TF in one cell type; the ten most strongly eNE-buffered TFs per cell type are outlined in blue. TFs in the upper-right quadrant (shaded blue) are buffered by both eNE and DREADD; those in the lower-left (shaded red) are amplified by both. c. Model. With aging, the dynamic range of activity-dependent transcription is compressed: induction fails for some genes, and the resting baseline rises toward the stimulated state for others. Activity nonetheless buffers part of the aging transcriptome in a cell type-specific manner.

We next asked whether direct activation of neurons could reproduce these same TF activities, using our chemogenetically activated (DREADD) dataset collected from female mice. Stimulating neurons by DREADDs recapitulated the buffering effect for the top TFs in neurons that were identified as buffered in eNE (**Fig. 6b, Extended Data Fig. 11a**). For example, both DREADDs and eNE activated CREB1 to oppose aging in hippocampal excitatory neurons and interneurons. When comparing DREADDs to an orthogonal aging state, defined in the physiological eNE dataset, the top eNE buffered and amplified TFs were also identified by DREADDs (**Extended Data Fig. 11b**). Although chemogenetic activation does not drive transcriptome-wide buffering (**Extended Data Fig. 10f**), these results suggest that it does reproduce buffering at the level of individual TFs and gene programs. Thus, the effects of novel experience on the aging transcriptome are at least partially attributable to the enhanced neuronal activity that accompanies novel experience.

Together, these results support a bidirectional model of the interaction between aging and activity in the brain. Aging compresses the dynamic range of youthful activity-regulated transcriptional programs. However, novel experience also stimulates TF activity and specific gene programs that can oppose, and in some cases amplify, the molecular changes that accompany aging (**Fig. 6c**). The balance between opposition and amplification likely depends on the aging state of the cell: at middle age, activity has a net effect of buffering the entire transcriptome, whereas this effect is diminished in older animals (**Fig. 5b,d**), highlighting middle age as a key intervention timepoint both for slowing dynamic range compression and harnessing buffering potential.

## Discussion

The brain must remain responsive to experience across a lifetime. Whether this capacity diminishes with age, and the extent to which neuronal activity itself protects against or exacerbates brain aging have remained unresolved. Our study combines new physiological and chemogenetic stimulation paradigms with high-resolution single-cell profiling to identify the principles that govern experience-dependent transcription during aging. Although the youthful ARG program contracts, aged cells hold a latent competence that, when engaged by experience, can directionally oppose the effects of aging. This competence is itself a target of aging. It is strongest at middle age and erodes with age in a cell-type- and sex-dependent manner. Together, these findings position activity-dependent transcription as both a mediator of acute circuit plasticity and a mechanism to recalibrate the transcriptomes of long-lived brain cells over a lifetime.

Whether ARG induction truly declines with age has remained unclear. Studies of individual immediate early genes have reported conflicting effects of age, including decreases, no change, and even increases^2,20,21^. Recent engram profiling found weaker recall-induced transcription in aged hippocampal neurons, most prominently among synaptic second-wave genes in DG granule cells^61^. Our data also shows that the dynamic range of the inducible transcriptome is compressed with age, and in males this compression converges on DG granule neurons. Behavioral paradigms such as novel experience and recall, however, cannot distinguish an aged animal that engages less from an aged cell that responds less. Here, using our chemogenetic activation paradigm, we find that under time-locked stimulation with matched drug exposure, aged neurons failed to mount the full youthful response. This places at least part of the defect inside the aged cell and suggests that restoring circuit drive alone in old brains may not be sufficient; the machinery that converts activity into transcription must also be preserved.

Our data point to two distinct routes by which aging could impair the normal youthful response. In failed induction, the stimulus arrives but the gene does not respond, implicating deficits in stimulus–transcription coupling that may be linked to epigenetic and genomic regulation. ARG induction depends on activity-regulated enhancers^62^, and learning-induced histone acetylation is lost in aged mice, a deficit whose rescue restores memory^63^. ARG loci are also sites of recurrent activity-induced DNA breaks^19,64,65^, and neuronal enhancers are hotspots of DNA repair^66,67^. As somatic mutations accumulate in aging neurons^68,69^, the loci most often used for plasticity may be among those most at risk of damage. In basal alteration, the gene is already altered at rest, leaving less room for further activity-dependent alteration. Basal alteration genes were enriched for the targets of inflammatory signaling transcription factors such as NF-κB, suggesting that increased inflammation during aging may occupy the dynamic range that activity would otherwise use^55^. However, because we sampled a single time point after stimulation, we cannot exclude that some apparent failures in ARG induction reflect delayed or truncated rather than absent induction. Time-resolved profiling, together with mapping chromatin and DNA damage at ARG loci in stimulated aged cells, will be needed to further understand the mechanisms that underly dynamic range compression.

The most unexpected finding of this study is that neuronal activity engages a restorative capacity in aged cells. Aged neurons recruited age-emergent synaptic programs absent in youth. Across the transcriptome, experience opposed basal aging in nearly every hippocampal cell type at middle age and, in aged males, returned it toward its youthful state. The aged response is therefore not a faded copy of the young one but a reorganized program that loses youthful components, gains new ones, and retains the capacity to counteract aging. These findings could shed light on why some cell types are more transcriptionally resilient to aging than others. Many aging atlases show that glia shift their transcriptomes more with age than neurons^27,28^. An intriguing possibility is that neurons are partly shielded because they engage activity-dependent programs directly and continually, renormalizing their transcriptome with each bout of activity. If so, the cognitive benefits of enrichment, exercise, and novelty in aged animals and humans^6,7,10,13^ may reflect a molecular maintenance function of experience. This function would act repeatedly against the slow drift of aging on the transcriptome.

Buffering, however, was not the only consequence of stimulation. Novel experience simultaneously amplified the activity of stress-responsive and hormonal regulators, including the glucocorticoid receptor, NF-κB and steroid hormone receptors. Amplification need not be harmful. Glucocorticoid signaling is integral to the activity-dependent response in glia^50^, and transient engagement of stress pathways can be hormetic, priming cells to withstand later challenges^70^. Amplification may therefore reflect the energetic and proteostatic cost of mounting a response in an aged cell, or an adaptive reinforcement of protective programs. Understanding how buffering and amplification are balanced within a cell is thus crucial for deciphering the overall effects on enhanced activity on aging cell function.

An important unresolved question is how neuronal activity promotes transcriptional buffering or amplification. One possibility is that activity reshapes chromatin itself, opening AP-1-bound enhancers in ways that persist after stimulation^42,71^. Another potential mechanism invokes competition for limiting cofactors. Activity-dependent transcription factors and inflammatory factors such as NF-κB draw on shared coactivators such as CBP/p300^72^, suggesting that activation could pull these resources away from inflammatory loci and thus reduce inflammatory gene expression in aged cells. However, our current identification of buffered TFs rest on curated regulon networks and remain correlative. Profiling CREB1 and ATF6 occupancy before and after stimulation, and acutely manipulating these factors during activation, will be needed to establish whether they causally shape buffering. Future experiments should establish how long buffering effects induced by novel experience last after a single exposure and whether repeated bouts of experience can sustain longer term changes.

It is noteworthy that direct chemogenetic activation of neurons reproduced aspects of buffering at the level of individual TFs but did not induce the effect on the transcriptome as a whole. These results suggest that activity may be beneficial, but that its pattern and dose matter. The full response appears to require the multicellular and systemic signals that accompany lived experience, including glial, vascular and hormonal inputs that a neuron-restricted stimulus cannot provide. This distinction is directly relevant to emerging therapies. Deep brain stimulation of memory circuits is under clinical evaluation in Alzheimer’s disease^73^, non-invasive approaches such as transcranial and sensory gamma stimulation are being tested in aging and dementia^74–76^, and chemogenetic tools are advancing toward the clinic^77^. Our findings raise the possibility that such interventions act in part by engaging the buffering capacity of stimulated cells. They also suggest that their efficacy will likely depend on whether stimulation reaches the relevant cell types and is delivered at an age where this capacity remains intact.

Vulnerability of the ARG program was shaped by cell type and sex. In males, deficits converged on DG granule neurons, a population repeatedly identified as an early casualty of aging^78,79^, while buffering capacity was largely retained into old age. In females, erosion of the youthful ARG program extended across neurons and glia, and buffering present at middle age had largely declined by old age. This results aligns with the observed cognitive decline reported in aged female mice^80,81^ and the higher prevalence of Alzheimer’s disease in women^82^. Steroid hormone receptor programs were among the regulators engaged by stimulation, raising the possibility that hormonal signaling tunes buffering capacity. Female mice, however, undergo reproductive senescence in middle age^83^ and may be biologically older at a given chronological age. Disentangling hormonal status from biological age will require dedicated designs that can carefully dissect the role of sex in ARG regulation during aging.

Our study recasts brain aging not as the loss of activity-dependent plasticity, but as its reorganization. Defining separable routes to failure suggests distinct intervention entry points: protecting chromatin and genome integrity to rescue induction and dampening inflammatory signaling to restore dynamic range. The persistence of a latent, activity-engaged capacity to oppose transcriptional aging offers a molecular rationale for the benefits of an engaged life and a framework for understanding the stimulation interventions that keep the aging brain responsive.

## Supporting information

Supplemental Table 1: Quality control measures for eNE and DREADD single-cell datasets

Supplemental Table 2: Differentially expressed genes across age and condition for all cell types

Supplemental Table 3: Buffered and amplified gene sets and transcription factor activities

## Methods

### Supplemental Tables

**Supplemental Table 1:** Quality control measures for eNE and DREADD single-cell datasets

**Supplemental Table 2:** Differentially expressed genes across age and condition for all cell types

**Supplemental Table 3:** Buffered and amplified gene sets and transcription factor activities

#### Contact for Resource and Material Sharing

Further information and requests for resources and reagents should be directed to and will be fulfilled by Elizabeth Pollina.

### Experimental Models and Analysis

#### Animals

Wild-type C57BL/6J mice of both sexes were obtained from the National Institute on Aging (NIA) aged rodent colony. For enhanced novel environment (eNE) experiments, young (4-month), middle-aged (14-month) and old (24-month) mice were used. For DREADD experiments, young (4-month) and old (20-month) mice were used. To mark neurons activated during novel environmental exposure, we used the *Fos-iCreERT2* (TRAP2) line (Jax# 030323)^84^ crossed to the *Ai14 tdTomato* reporter (Jax # 007914)^85^. On arrival, cage-mates were housed together and given at least two weeks to acclimate to the facility before experiments began, minimizing neuronal activity caused by environmental novelty. Within each sex, animals from all age groups were processed on the same day, and males and females were collected on successive days within one week. Mice were housed under standard conditions maintained by the Washington University Division of Comparative Medicine, with food and water available ad libitum. All procedures were approved by the Washington University Institutional Animal Care and Use Committee (protocol #25-0326).

#### Novel environment exposure

To induce physiological neuronal activity *in vivo*, wild-type C57BL/6J mice (n = 3 per condition per age group per sex; 36 mice total) were exposed to a custom novel enriched environment for 6 hours. We collected males and females on separate days within the same week to minimize the effects of pheromones during the eNE paradigm; thus, we report the aging biology observed within each sex but do not formally test for sex-dependent effects. Mice were removed from their standard home cages (where they were housed in groups of five) and placed into three enrichment enclosures, one per age group, with three age-matched mice per cage. The enrichment enclosure was a custom two-compartment structure. The first compartment was divided into two sections each containing food and water, with the food side additionally equipped with two running wheels, plastic tubes, ramps, assorted enrichment objects, and a red plastic house with four entrances. The second floor was accessible via ramps and featured a maze constructed from interlocking blocks. For comparison with a conventional paradigm (**Fig. 1c**), a separate group of mice was exposed for 6 hours to a classical novel environment consisting of a standard single-level rat cage containing a tunnel, a horizontal running wheel and assorted enrichment objects. Home-cage control (HC) mice of matched age and sex remained undisturbed in their standard home cages for the equivalent 6-hour duration. Active engagement with the novel environment was confirmed by continuous video surveillance throughout the exposure period. Body weight was recorded for each animal immediately following exposure. To ensure that exposure duration did not exceed 6 hours, eNE and HC mice were dissected in a staggered fashion within the 6-hour window, with the order across age groups rotated each experimental round to eliminate order bias. All mice were transcardially perfused with 30 mL of ice-cold 1× phosphate-buffered saline (PBS) to remove circulating blood.

#### Activity-dependent labeling with *Fos-TRAP2*

To mark neurons activated during eNE exposure, we used the *Fos-iCreERT2* (TRAP2) crossed to the *Ai14 tdTomato* reporter, with Cre-negative littermates as recombination controls. CreERT2 is expressed from the endogenous *Fos* locus but remains cytoplasmic until 4-hydroxytamoxifen (4-OHT) permits its nuclear entry. Thus, tdTomato recombination marks only neurons in which *Fos* is induced during the 4-OHT window. To minimize activity-dependent labeling caused by handling and injection stress, mice were handled daily and given mock intraperitoneal injections of 1× PBS for 2 weeks before the experiment, and body weight and food consumption were recorded over this period.

4-OHT (Sigma-Aldrich, H7904) was dissolved in ethanol at 22.2 mg/mL by heating for 5 min at 37 °C, mixed with an equal volume of Chen oil (1:4 castor oil:sunflower oil), vortexed and heated for 5 min at 37 °C, and the ethanol was evaporated by vacuum centrifugation for at least 1 h; the residue was then diluted with Chen oil to a final concentration of approximately 6.7 mg/mL. The solution was prepared fresh and used on the day of injection. On the day of the experiment, 4-month-old mice received a single intraperitoneal injection of 4-OHT (50 mg/kg; 8 µL per g body weight) immediately before being placed in the enhanced novel environment (eNE) for 6 hours, after which they were returned to their home cages. Home-cage controls were injected on the same schedule and remained in their home cages throughout. Males and females were run on consecutive days. Mice were perfused 7 days after 4-OHT to allow tdTomato expression. Brains were immersion-fixed in 4% PFA overnight at 4 °C, cryoprotected in 20% sucrose, and cut into free-floating sections (30 µm); native tdTomato fluorescence was imaged without antibody amplification. A single cohort was used, comprising CreERT2-positive mice of both sexes (new eNE, n = 5; home cage, n = 6); representative image is shown in **Extended Data Fig. 1b**.

#### Tissue collection

Following perfusion, the left hemisphere was immersion-fixed in 4% paraformaldehyde (PFA) overnight at 4 °C, cryoprotected in 30% sucrose and embedded in OCT compound. It was then cryosectioned at 30 µm for free-floating immunohistochemistry or at 20 µm for RNAscope. The right hippocampus was microdissected, snap-frozen in liquid nitrogen and stored at −80 °C for single-cell RNA sequencing. Before experiments, mice were assessed for general health, and animals that were visibly underweight or unresponsive to handling were excluded. At perfusion and dissection, the abdominal organs and brain were inspected, and animals with gross abnormalities such as tumors were excluded from single-cell sequencing.

#### Immunostaining and RNAscope

Fixed-frozen coronal sections (20 µm) mounted on SuperFrost Plus slides were dried for at least 30 min, outlined with a hydrophobic barrier pen and washed twice for 10 min in 1× PBS. Antigen retrieval was performed by steaming slides in citric acid buffer (10 mM citric acid, 0.05% Tween-20, pH 6.0) at 100 °C for 10 min, followed by washes in 1× PBS. Sections were blocked for 1 h at room temperature in 5% normal goat serum with 0.3% Triton X-100 in 1× PBS, then incubated overnight at 4 °C with primary antibodies against FOS (Synaptic Systems, cat. no. 226-308; 1:500) and NPAS4 (in-house; 1:1,000) diluted in blocking buffer. Slides were washed three times for 10 min in 1× PBS containing 0.1% Triton X-100 followed by 1× PBS, incubated for 1–2 h at room temperature with an Alexa Fluor 647-conjugated goat anti-guinea pig secondary antibody (Invitrogen, cat. no. A21450; 1:500) and a goat anti-rabbit secondary antibody (Invitrogen, cat. no. A32731; 1:500) together with DAPI (1:1,000) in 1× PBS, and washed again as above. Coverslips were mounted with Fluoromount-G and sections were imaged on a Leica THUNDER Imager (inverted DMi8).

*In situ* hybridization was performed using the RNAscope Multiplex Fluorescent Reagent Kit v2 (Advanced Cell Diagnostics) on fixed-frozen tissue. Mice were perfused with 1× PBS, and brains were post-fixed in 4% paraformaldehyde (PFA) in 1× PBS overnight at 4 °C, cryoprotected in 30% sucrose in 1× PBS until equilibrated, and stored at −80 °C. Coronal cryosections (20 µm) were mounted on SuperFrost Plus slides, dried for 5–10 min at room temperature, washed in 1× PBS, baked at 60 °C for 30 min and post-fixed in prechilled 4% PFA for 15 min at 4 °C. Sections were dehydrated through a graded ethanol series (50%, 70%, 100%, 100%) and treated with RNAscope Hydrogen Peroxide for 10 min at room temperature, followed by Protease III for 30 min at 40 °C. Probes targeting *Npas4* (RNAscope Probe-Mm-Npas4-C2, ACD, cat. no. 423431-C2) and *Fos* (RNAscope Probe-Mm-Fos-C3, ACD, cat. no. 316921-C3) were hybridized for 2 h at 40 °C, followed by sequential amplification steps (Amp 1–3) and HRP-based signal development with TSA Plus fluorophores diluted 1:1,000 in TSA buffer. These sections were counterstained with DAPI, mounted with the provided mounting reagent and imaged on a Zeiss LSM 880 confocal microscope. In a separate set of sections, *Adm* was detected with RNAscope Probe-Mm-Adm-C1 (ACD, cat. no. 493601) using the same protocol, followed by immunostaining for GFAP to identify astrocytes. GFAP was detected with a guinea pig recombinant monoclonal anti-GFAP antibody (clone Gp134B1, Synaptic Systems, cat. no. 173 308; 1:500) and an Alexa Fluor 647-conjugated goat anti-guinea pig IgG (H+L) highly cross-adsorbed secondary antibody (Invitrogen, cat. no. A-21450; 1:500). These sections were counterstained with Hoechst and imaged on a Leica THUNDER Imager (inverted DMi8).

#### Quantification of RNAscope and immunostaining

FOS and NPAS4 signals were quantified in Fiji (ImageJ) using custom macros. Regions of interest for the dentate gyrus, CA1, CA2 and CA3 were drawn manually on the Hoechst reference channel. Each channel was converted to 8-bit, smoothed with a Gaussian blur (σ = 1 for target channels, σ = 4 for the reference) and binarized with the Triangle auto-threshold, excluding background pixels from the threshold calculation for target channels. Signal was expressed as the thresholded target area divided by the thresholded Hoechst area within each region. Immunostained sections were imaged with FOS in the 647 channel and NPAS4 in the 568 channel; RNAscope sections with *Fos* in the 570 channel and *Npas4* in the 520 channel. The same parameters were applied to all images.

*Adm* expression in astrocytes was quantified in Imaris 10.1.1 (Oxford Instruments). Z-stacks acquired with a 20× objective were processed with Large Volume Computational Clearing and collapsed into maximum intensity projections. Nuclei were detected in the Hoechst channel using the Spots function (estimated diameter 15 µm, background subtraction enabled) and filtered by quality. Spots with GFAP intensity above a fixed threshold were defined as astrocytes, and astrocytes with *Adm* intensity above a fixed threshold as *Adm*-positive. The same detection and classification parameters were applied to all images.

#### Stereotaxic viral injection

Young (4-month) and old (20-month) male and female C57BL/6J mice were anesthetized with isoflurane (induction 2–4%, maintenance 1.0–1.9% at <200 mL/min) and mounted on a stereotaxic apparatus. Buprenorphine SR (1 mg/kg, subcutaneous) was administered for perioperative analgesia. Following a midline scalp incision and skull exposure, the head was leveled using bregma–lambda Z-axis calibration (tolerance <0.05 mm) and bilateral coordinate symmetry was confirmed (lateral Z-axis tolerance <0.05 mm). AAV9-hSyn-hM3Dq-T2A-mScarlet-KASH was produced by the Hope Center Viral Vectors Core (Washington University in St. Louis) and diluted to a working titer of 1.0×10¹² Gc/mL. Each mouse received bilateral hippocampal injections at four sites: dorsal hippocampus (ML ±1.5 mm, AP −2.0 mm, DV −1.5 mm from bregma) and ventral hippocampus (ML ±2.88 mm, AP −3.10 mm, DV −2.3 mm from bregma), delivering 1 µL per site (4 µL total per mouse) at a controlled rate. The needle was held in place for 5–10 minutes following each injection before slow retraction to minimize reflux. The scalp was closed with interrupted sutures and veterinary tissue adhesive. Mice were recovered on a heated pad and monitored at 15-minute intervals for a minimum of 30 minutes. Viral expression was allowed to proceed for 3 weeks prior to chemogenetic activation.

#### Chemogenetic activation and tissue collection

Agonist C21 (C21; Hello Bio, HB6124; 2 mg/kg, intraperitoneal) was administered to mice exactly 2 hours prior to perfusion. Control mice received weight-equivalent volumes of PBS. All animals were transcardially perfused with 30 mL of ice-cold 1× PBS at the 2-hour timepoint. Following perfusion, the left cerebral hemisphere was immersion-fixed in 4% PFA overnight at 4°C and cryoprotected in 20% sucrose for histological validation of viral expression and mScarlet distribution. The right hemisphere was used for molecular analyses: the hippocampus was microdissected and bisected into dorsal and ventral portions, and the cortex was divided into dorsal, middle, and ventral thirds for analysis of C21 exposure by LC-MS (see below). All tissues were immediately frozen in liquid nitrogen and stored at −80°C until processing.

#### C21 penetrance validation by LC-MS

Because the hippocampus was reserved for scRNA-seq, we quantified C21 in the cortex from the same animals as a surrogate for brain exposure. We first established by mass spectrometry (MS) that C21 is equivalent between cortical and hippocampal matrices when dissected from the same mice (**Extended Data Fig. 1d**). To confirm central nervous system (CNS) penetrance of C21 at the 2-hour timepoint, cortical and hippocampal tissue from a subset of injected animals was processed for LC-MS/MS. Brain tissue was lysed by tip sonication in methanol, and 30 mg of lysate was subjected to target extraction using a MeOH:MTBE mixture (3:10, v/v) with vortex mixing for 1 minute followed by overnight incubation at −80°C. Following centrifugation, the hydrophilic fraction was collected, dried by SpeedVac without heat, and reconstituted in 100 µL of 50% methanol. Chromatographic separation was performed on a Waters ACQUITY Premier HSS T3 column (1.7 µm, 2.1 mm × 150 mm) using a 10-minute gradient with mobile phase A (0.1% formic acid in water) and mobile phase B (0.1% formic acid in 90% acetonitrile), with 2 µL injected per run. C21 was detected in positive ionization mode on an Orbitrap ID-X Tribrid mass spectrometer coupled to a Vanquish Horizon UHPLC system (Thermo Scientific), with MS2 ions quantified by parallel reaction monitoring (PRM). Collision energy was optimized by stepwise increment. Fragment ion peak areas were quantified using Skyline 25.1. Calibration curves were generated by serial dilution of C21 light standard across 8 concentration levels (2.4 ng/mL to 2.5 µg/mL) in both cortical and hippocampal matrix, yielding linear dynamic ranges with R² = 0.987 (cortex) and R² = 0.990 (hippocampus). As no isotope-labeled internal standard was used, C21 concentrations are reported as semi-quantitative estimates based on peak area. Relative comparisons between samples were performed on raw peak areas. C21 was detectable and quantifiable in both cortex and hippocampus, confirming adequate CNS penetrance at the 2-hour collection timepoint.

#### Histological validation and quantification of viral expression

To confirm viral transduction and characterize mScarlet expression, one anterior coronal hippocampal section per animal was imaged on a Zeiss LSM 880 confocal microscope in four channels: FOS (647 nm), mScarlet (561 nm), NPAS4 (488 nm) and DAPI. Images were analyzed in Fiji. A fixed threshold was set for each channel on the 16-bit images (identical across all images; FOS 580, mScarlet 12,300, NPAS4 5,027 and DAPI 4,670 gray levels). The images were then converted to 8-bit and to binary masks, and the channels were split. Regions of interest (ROIs) were drawn manually on the DAPI channel around the principal cell layers of the dentate gyrus (granule cell layer) and of CA3, CA2 and CA1 (pyramidal cell layer), where mScarlet signal was concentrated. For each ROI, the area and mean gray value of the binary mask were measured in each channel; in a binary mask the mean gray value scales with the fraction of the ROI above threshold. Viral expression was quantified with two measures (**Extended Data Fig. 1,8a**). First, mScarlet signal was normalized to nuclear density as the ratio of mScarlet to DAPI mean gray value within each ROI (mScarlet⁺/DAPI⁺ area ratio). Second, viral spread was quantified as the percentage of ROI area above the mScarlet threshold (mScarlet⁺ area, % of ROI). Each measure was averaged across the four subregions (dentate gyrus, CA3, CA2 and CA1) to give one value per animal. Males and females were analyzed separately (females: young C21 n = 5, young PBS n = 4, old C21 n = 4, old PBS n = 3; males: young C21 n = 5, young PBS n = 4, old C21 n = 3, old PBS n = 3).

#### Validation of DREADD-induced neuronal activity

Neuronal activity induced by DREADD was validated by immunohistochemistry for the immediate early gene protein products FOS and NPAS4. Brains were sectioned coronally at 30 µm on a cryostat, and free-floating sections were incubated with primary antibodies against FOS (Synaptic Systems, cat. no. 226-308, 1:500) and NPAS4 (in-house, 1:1000), followed by species-appropriate secondary antibodies (anti-guinea pig, Invitrogen cat. no. A21450, 1:500; anti-rabbit, Invitrogen cat. no. A32731, 1:500). Sections were imaged on a Zeiss LSM 880 confocal microscope in four channels: 647 nm (FOS), 488 nm (NPAS4), 561 nm (mScarlet), and DAPI. Images were converted from 16-bit to 8-bit, and a manual intensity threshold was applied uniformly across all images within each channel prior to channel splitting. Regions of interest were defined by DAPI signal and subdivided into CA1, CA2, CA3, and dentate gyrus subregions. Mean gray intensity was measured per channel and normalized to DAPI to control for sectioning variability. DREADD activation induced a significant increase in FOS immunoreactivity across hippocampal subregions relative to PBS-treated controls (approximately 7–10-fold enrichment).

#### Single-Cell RNA-Sequencing

##### Tissue preparation and nuclei isolation

The right hippocampus was dissected immediately after cardiac perfusion and snap-frozen in liquid nitrogen. Fixed cells were prepared from frozen tissue according to the 10x Genomics Chromium Fixed RNA Profiling sample preparation protocol, then counted and quality-assessed before library construction.

##### Library preparation and sequencing

eNE libraries were constructed with Chromium Fixed RNA Profiling (Flex) chemistry using the Chromium Mouse Transcriptome Probe Set v1.1.1 (10x Genomics). DREADD libraries were constructed with GEM-X Flex v2 chemistry using the Chromium Mouse Transcriptome Probe Set v2.0.0, supplemented with three custom probe pairs for each of mScarlet and hM3Dq to detect virally expressed transcripts within the same assay. Each pair consisted of a left-hand (LHS) and a right-hand (RHS) probe designed according to 10x Genomics Flex custom probe guidelines (sequences 5′→3′):

**mScarlet probe 1:**

LHS CCTTGGCACCCGAGAATTCCAAGCGTTCGTACTGTTCCACCACGGT; RHS GTAGTCCTCGTTGTGGGAGGTGATGCCCATATAAGAAA.

**mScarlet probe 2:**

LHS CCTTGGCACCCGAGAATTCCACGATCTCGAACTCGTGGCCGTTCAT; RHS GGAGCCCTCCATGTGCACCTTGAACCCCATATAAGAAA.

**mScarlet probe 3:**

LHS CCTTGGCACCCGAGAATTCCAAAGTTGGTGCCGCGGAGCTTCACCT; RHS TGTAGATCAGGGTGCCGTCCTCCAGCCCATATAAGAAA.

**hM3Dq probe 1:**

LHS CCTTGGCACCCGAGAATTCCATGTCTCTGCCTCTGTCCCAGAGGCT; RHS TGCAGGCCAGCAAGCTCTTTGGTACCCCATATAAGAAA.

**hM3Dq probe 2:**

LHS CCTTGGCACCCGAGAATTCCACCAGGGAGGCAGCAGCATCATTGTT; RHS GTTCCAACTGTCACTGCTGCTGTGGCCCATATAAGAAA.

**hM3Dq probe 3:**

LHS CCTTGGCACCCGAGAATTCCACTTAGCTGTGTCCACGGCTGACTCT; RHS AGCTGGATGGGAAGCTTGGAGAAGCCCCATATAAGAAA.

Because the hM3Dq probes showed low capture, only mScarlet was used in downstream analyses. Libraries were sequenced on an Illumina NovaSeq X Plus at the Genome Technology Access Center at the McDonnell Genome Institute (GTAC@MGI, Washington University in St. Louis) across four runs, at a target depth of ∼400 million read pairs per sample. eNE samples comprised 16 male samples (SR008865), 16 female samples (SR009556) and four middle-aged home-cage samples sequenced in a supplemental run (SR011142). DREADD samples comprised 12 female samples (SR011397).

##### Read alignment and quantification

Reads were demultiplexed and aligned to the mouse reference genome (GRCm39-2024-A) using Cell Ranger (10x Genomics) v9.0.1 for eNE samples and v10.0.0 for DREADD samples. Filtered feature-barcode matrices were imported into Seurat (v5.5.0) for downstream analysis.

##### Quality control and filtering

Per-sample quality control was performed in Seurat (v5.5.0), applying identical criteria to the eNE and DREADD datasets. For each sample, cells were retained if their number of detected features fell between the 25th percentile and the 75th percentile plus 1.5× the interquartile range of that sample’s distribution, if they had at least 100 unique molecular identifiers (UMIs), and if less than 10% of their reads came from mitochondrial transcripts. Ambient RNA contamination was removed using SoupX, and doublets were removed using scDblFinder. In total, 818,098 cells from 36 eNE samples and 322,130 cells from 12 DREADD samples passed quality control (**Supplemental Table 1**).

##### Normalization, dimensionality reduction, and clustering

The eNE and DREADD datasets were processed and integrated separately using the same pipeline. Filtered count matrices were normalized using the LogNormalize method (scale factor = 10,000), and the top 2,000 highly variable features were identified with the vst method. Samples were merged and split into layers by experimental group (sex × age for eNE; age × treatment for DREADD), and 10,000 cells were sketched from each group (60,000 eNE and 40,000 DREADD cells). Principal component analysis (50 components) was performed on the sketched cells, and layers were integrated using Seurat’s reciprocal PCA (RPCA) method. A shared nearest-neighbor graph (k = 20) was constructed on the first 30 integrated dimensions, and sketched cells were clustered using the Louvain algorithm at a resolution of 0.3. All cells were then projected onto the integrated space and sketched cluster labels. In the eNE dataset, no cluster was specific to one sex: the proportion of female cells per cluster (median 0.44; interquartile range 0.42–0.46) closely matched the dataset-wide proportion. Clusters in both datasets were annotated manually using canonical marker genes, as described below.

##### Cell Type Annotation

Unsupervised clusters were annotated manually based on the expression of curated canonical marker genes visualized as stacked violin plots. Markers were organized hierarchically into broad and cell-type-specific groups. Broad neuronal identity was confirmed using pan-excitatory markers (*Slc17a7*, *Slc17a6*, *Cux2*, *Satb2*, *Neurod2*, *Sv2c*, *Car10*, *Ntng1*, *Tbr1*, *Tyro3*, *Foxp2*), pan-neuronal markers (*Rbfox3*, *Map2*, *Sv2b*) and neural stem cell markers (*Sox2*, *Nes*, *Fabp7*, *Crip2*, *Aldoc*, *Mt3*, *Dbi*). Hippocampal excitatory neuron subtypes were distinguished using subregion-specific markers: dentate gyrus neurons (*Prox1*, *Glis3*, *Egr3*, *C1ql2*, *C1ql3*, *Itpka*), CA3 neurons (*Cpne4*, *Spock1*, *Lhx9*, *Neurod6*, *Coch*), CA1 neurons (*Spink8*, *Mpped1*, *Zfhx4*, *Cacng5*, *Wfs1*, *Fibcd1*, *Lefty1*), CA2 neurons (*Amigo2*, *Pcp4*, *Rgs14*, *Zfp536*, *Ovol2*, *Fezf2*, *Map3k15*, *Tiam2*) and subicular neurons (*Tshz2*, *S100b*, *Dlk1*, *Tpbg*, *Fn1*, *Gpc3*, *Cbln4*, *Col5a2*, *Ly6g6e*). Inhibitory interneurons were identified by expression of *Gad1*, *Gad2*, *Slc32a1*, *Pvalb*, *Sst*, *Vip* and *Kit*. Glial populations were annotated using astrocyte markers (*Gfap*, *Aldh1l1*, *Aqp4*), oligodendrocyte markers (*Olig1*, *Olig2*, *Ptgds*, *Mag*, *Mog*), OPC markers (*Pdgfra*) and microglia markers (*C1qc*, *Itgam*, *Trem2*, *Cx3cr1*). Non-neural populations were identified using markers for endothelial cells (*Pecam1*, *Flt1*, *Cldn5*), pericytes (*Pdgfrb*, *Kcnj8*, *Abcc9*), astrocyte precursor cells (APCs; *S100a6*), stromal cells (*Col1a1*, *Col1a2*, *Dcn*), Cajal–Retzius cells (*Reln*, *Trp73*, *Lhx1*), macrophages (*Cd68*, *Csf1r*, *Ms4a7*), epithelial cells (*Epcam*, *Krt8*, *Krt18*) and T cells (*Cd3e*, *Cd3d*, *Cd247*). Normalized expression values were retrieved from the RNA assay data layer using Seurat’s FetchData function (see **Extended Data Fig. 3**)

#### Single-cell activity scores

As an independent estimate of neuronal activation, per-cell activity scores were computed with NEUROeSTIMator^51^ (neuroestimator R package, species = “mmusculus”), a deep-learning model trained on immediate-early gene expression. Raw UMI counts for all cells were scored in batches of 50,000, returning one score between 0 and 1 per cell. Scores were averaged across cells within each neuronal cell type (DG, CA1, CA2, CA3 and subicular excitatory neurons, and interneurons), sex, age and condition, and are shown as the mean ± SEM across cells (**Fig. 2c**).

#### Subclustering analysis for astrocytes and inhibitory neurons (INs)

Inhibitory neurons (61,890 cells) and astrocytes (103,148 cells) from the eNE dataset were extracted and reanalyzed separately. For interneurons, counts were log-normalized, the top 2,000 highly variable features were scaled, and principal component analysis was performed (30 components). Principal components were integrated across samples with Harmony (v2.0.5), and a shared nearest-neighbor graph was constructed on the first 20 Harmony dimensions. Cells were clustered using the Louvain algorithm across resolutions from 0.5 to 0.8, and a resolution of 0.55 (20 clusters) was selected. Each cluster was assigned to a subtype by manual inspection of canonical marker expression in dot plots and feature plots, supported by Seurat’s FindAllMarkers (Wilcoxon rank-sum test; min.pct = 0.25, log2 fold-change threshold = 0.25). Markers were PV (*Pvalb*, *Kcnc1*, *Kcnmb4*, *Pthlh*, *Vipr2*), SST (*Sst*, *Npy*, *Cort*, *Chodl*, *Hpse*, *Elfn1*), VIP (*Vip*, *Calb2*, *Cck*, *Chrm2*, *Rxrg*) and LAMP5 (*Lamp5*, *Nos1*, *Ndnf*, *Id2*), together with pan-interneuron (*Gad1*, *Gad2*, *Slc32a1*) and developmental origin (MGE: *Lhx6*, *Nkx2-1*; CGE: *Sp8*, *Etv1*) markers. Clusters were assigned to PV (5 clusters), SST (6), VIP (4) and LAMP5 (4) subtypes. One cluster without a clear subtype signature was labeled ambiguous and excluded from subtype analyses.

For astrocytes, counts were log-normalized, the top 3,000 highly variable features were identified, and principal component analysis was performed (30 components). A shared nearest-neighbor graph was constructed on the first 20 principal components, and cells were clustered using the Louvain algorithm at a resolution of 0.1. This yielded five clusters (C1–C5; 77,680, 12,820, 9,420, 2,316 and 912 cells). Clusters were characterized using *Gfap*, *Agt*, *Ogt*, *Fam107a*, *Frzb*, *Ascl1*, *Aldh1l1*, *S100b*, *Vim*, *Hopx*, *Id3* and *Mfge8*.

Differential expression between eNE and home-cage conditions was tested within each subtype at each age using pseudobulk DESeq2. Counts were summed per animal, genes with at least five counts in at least three samples were retained, and comparisons were run only when each condition had at least three animals. Sexes were analyzed separately (design ∼ condition). Genes with padj < 0.05 were considered significant. Subtype-specific ARGs were defined as genes significantly induced in one subtype but in no excitatory neuron population and no other subtype of the same class, identified independently at each age (see **Extended Data Fig. 5e, Extended Data Fig. 6d**).

#### eNE Pseudobulk differential expression analysis

Differential gene expression was assessed using a pseudobulk approach. For each cell type, raw UMI counts were summed across cells within each biological replicate (individual animal) using the AggregateExpression function in Seurat v5, grouping by cell type, sample identity, experimental condition, age and sex. Within each comparison, genes were retained if they had at least 10 counts in at least two pseudobulk samples. Comparisons with fewer than two samples in either group were excluded; as a result, not all cell types are included in every sex and age analysis.

DESeq2 was used to model differential expression with a two-group design (∼ group), using default settings and Wald tests; log_2_ fold changes (log_2_FC) were not shrunk. Analyses were performed separately within each sex, and the sexes were not compared directly because they were sequenced in separate batches. The following pairwise comparisons were performed: enhanced novel environment versus home cage within each age group (young, middle-aged, and old); age-associated changes within home-cage animals (old vs. young, old vs. middle-aged and middle-aged vs. young); and age-associated changes within eNE-exposed animals (the same three contrasts). For the buffering index, two cross-condition, cross-age comparisons were performed: old eNE versus young home cage, and middle-aged eNE versus young home cage. Statistical significance was defined as Benjamini–Hochberg adjusted p < 0.05.

#### DREADD pseudobulk differential expression analysis

For the DREADD dataset, neuronal cells were pre-filtered to retain only those with ≥ 50 raw mScarlet UMI counts. This threshold corresponds to the 99th percentile of mScarlet counts in non-transduced glial cells and excludes neurons whose mScarlet signal likely reflects ambient RNA contamination. All glial cells were retained regardless of mScarlet expression. Interneuron subtypes (IN1 and IN2) were merged into a single interneuron class (IN) prior to analysis. Raw UMI counts were aggregated per cell type per animal using AggregateExpression in Seurat v5. Genes were retained if they had ≥ 10 counts in ≥ 2 pseudobulk samples, and comparisons with fewer than four pseudobulk samples in total, or fewer than two in either group, were excluded.

We used two approaches to account for differences in mScarlet expression: a covariate approach, which included median mScarlet expression per mouse as a covariate in the model, and a downsampling approach, which equalized mScarlet levels and cell numbers across samples. The covariate approach retains more cells and therefore samples a larger pool of responses, whereas the downsampling approach provides a stringent match for DREADD expression. In the covariate approach, the per-mouse, per-cell-type median mScarlet count (scaled to unit variance) was included as a continuous covariate in the DESeq2 model for neuronal cell types (design: ∼ mScarlet_median_scaled + group); glial cell types were modeled without this covariate (design: ∼ group). In the downsampling approach, per-mouse 10th–90th percentile mScarlet bounds were computed for each neuronal cell type, and the median bounds across mice were applied as a common inclusion window; cells outside this window were excluded. All cell types were then downsampled so that each mouse × group combination contained the minimum observed cell count for that cell type (random seed = 42), and both neurons and glia were modeled with a two-group design (design: ∼ group) without an mScarlet covariate. We focus on the downsampling approach as the more stringent normalization and provide covariate results in Extended Data.

The following pairwise comparisons were performed under both approaches: C21 versus PBS within young animals; C21 versus PBS within old animals; aging-associated changes within PBS-treated animals (old vs. young); aging-associated changes within C21-treated animals (old vs. young); and old C21 versus young PBS, capturing the net transcriptional position of chemogenetically stimulated aged neurons relative to unstimulated young animals. Statistical significance was defined as Benjamini–Hochberg adjusted p < 0.05 in both approaches.

#### Gene classifications

##### Maintained, Lost, Age-emergent ARGs

To characterize how the activity-regulated gene (ARG) program identified in young mice is preserved or altered with aging, each DEG from the young eNE-versus-HC comparison (YeNE vs. YHC, padj < 0.05) was classified according to its regulation in the corresponding middle-aged (MeNE vs. MHC) and old (OeNE vs. OHC) comparisons. Classification was performed independently for each cell type, sex and age, with young ARGs serving as the anchor. Young-induced ARGs that remained significantly upregulated at the older age were classified as Maintained Upregulated, and young-suppressed ARGs that remained significantly downregulated were classified as Maintained Downregulated. ARGs that were significantly regulated in the opposite direction at the older age were classified as Flipped. Young-induced ARGs that were no longer significantly differentially expressed at the older age (padj ≥ 0.05) were classified as Lost Upregulated, and the equivalent young-suppressed ARGs were classified as Lost Downregulated. Genes significant in MeNE vs. MHC or OeNE vs. OHC but not significant or not detected in YeNE vs. YHC were classified as Age-emergent Up or Age-emergent Down. All comparisons used a Benjamini–Hochberg adjusted p cutoff of 0.05.

##### Failed induction versus basal alteration

Lost genes were further subdivided to distinguish two explanations. For each cell type, sex and age, the basal change with age was taken from the matched home-cage comparison (MHC vs. YHC for middle age; OHC vs. YHC for old age). For each lost gene, a basal-aging fraction was computed as its home-cage aging log_2_FC divided by its activity log_2_FC (*i.e.* YeNE vs. YHC). A lost gene was classified as basal alteration if this fraction was ≥ 0.5; that is, its basal expression had already shifted with age in the direction that eNE induces in young mice, by at least half the young eNE response. No significance threshold was applied to the basal age change. All other Lost Upregulated genes that did not meet this basal age change ≥ 0.5 were classified as failed induction, and all other Lost Downregulated genes as failed repression.

#### Pathway enrichment

Gene Ontology Biological Process enrichment was performed with clusterProfiler (enrichGO; org.Mm.eg.db) for each gene set per cell type, per cell class (neurons, glia and vascular cells), and pooled across cell types. Terms were considered significant at BH-adjusted p < 0.05 and q < 0.2. Gene sets with fewer than five genes were not tested, and redundant terms were merged with simplify (similarity cutoff 0.7). The same procedure was used for all pathway analyses in this study (eNE and DREADD); only the background differed by gene set. For ARG sets, the background was all genes in the anchor DESeq2 table (YeNE vs. YHC), pooled across cell types. For buffering gene sets, the background consisted of the genes eligible for that set (genes tested in the corresponding held-out contrasts), so that enriched terms reflected buffering rather than aging in general.

#### Dynamic range compression of the ARG Program

For each cell type and sex, genes significantly up- or downregulated in YeNE vs. YHC (padj < 0.05) were treated as separate anchor sets, and their log_2_FC values were extracted from YeNE vs. YHC, MeNE vs. MHC and OeNE vs. OHC regardless of significance at older ages. Genes without an estimate at a given age were excluded at that age, and cell types were analyzed if at least one anchor set contained ≥ 10 genes. Within the up- and downregulated sets separately, shifts in log_2_FC between ages were tested with two-sided paired Wilcoxon signed-rank tests on genes with estimates at both ages (≥ 5 genes). p values were Benjamini–Hochberg corrected across cell types and age comparisons within each sex and gene set (**Fig. 3b, Extended Data Fig. 7a,b**).

Compression was quantified with a compression score. For each set containing ≥ 10 genes with estimates at both ages, the score was the mean |log2FC| in YeNE vs. YHC divided by the mean |log2FC| at the older age over the same genes (capped at 10). Scores were averaged across the up- and downregulated sets, so values > 1 indicate a weaker eNE response with age. Scores were computed for the young-to-middle and young-to-old transitions. 95% confidence intervals and p values (the proportion of scores ≤ 1) were obtained from 1,000 bootstrap resamples of genes within each set. p values were Benjamini–Hochberg corrected across cell types within each sex and age (**Extended Data Fig. 7c**).

#### Conserved dynamic range compression in eNE and DREADD

To test whether compression of the activity response was shared between paradigms, DESeq2 results from female mice were compared between eNE (YeNE vs. YHC and OeNE vs. OHC) and DREADD (young C21 vs. young PBS and old C21 vs. old PBS; downsampling approach) (**Fig. 4g**). For each cell type, shared young ARGs were genes induced in both young contrasts (padj < 0.05). For each shared gene and paradigm, the retained response was calculated as r = log_2_FC_old / log_2_FC_young, and genes with r < 0.5 in both paradigms were classified as compressed in both. Each gene compressed in both paradigms was then given one value per age, its log**2**FC averaged across the cell types in which it was compressed in both. **Fig. 4g** shows the median and interquartile range of these values for all such genes (left), together with the 20 genes with the strongest compression (right), ranked by the smaller of their two young responses multiplied by the smaller of their two fractional declines. eNE middle-age responses (MeNE vs. MHC) are shown for reference only.

#### Transcriptional buffering analysis

##### Definitions and comparisons

Transcriptional buffering was defined as eNE-induced expression changes that opposed basal aging-associated changes at a given age. For each cell type, sex and age, the aging vector **B** contained the per-gene log_2_FC of middle-aged or old HC versus young HC mice, and the stimulation vector **E** contained the per-gene log_2_FC of eNE versus HC mice at the same age. Middle and old age were analyzed separately with age-matched comparisons. Each sex was analyzed on its own, and aging and stimulation vectors were never paired across sexes.

##### Held-out cosine design

Because B and E share the aged HC group with opposite signs, sampling noise in those mice enters both vectors and biases them toward opposition even when eNE has no effect. To remove this coupling, we used a held-out design. Each of the three aged HC mice was held out in turn (k = 1–3). The aging contrast compared the remaining two aged HC mice with the young HC mice (Bₖ), and the stimulation contrast compared the eNE mice with the held-out HC mouse (Eₖ), so no mouse contributed to both contrasts within an iteration. Each contrast was analyzed with DESeq2 using the same pseudobulk counts, gene filter and design (∼ group) as the main differential expression analysis, with Wald tests and unshrunken log2FC. Where standard dispersion estimation failed, gene-wise dispersion estimates were used. Each held-out iteration therefore required at least three aged HC mice, so that one could be held out for the stimulation contrast while two remained for the aging contrast, and at least three aged eNE mice. At least two young HC mice were required, as these enter the aging contrast only, where they serve as the reference group. The three held-out iterations were used for all cell-type, gene and transcription factor buffering analyses described below.

For each iteration, the cosine similarity between **B**ₖ and **E**ₖ was computed across genes with estimates in both (≥ 20 genes):

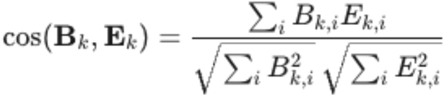

The three values were averaged to give the final held-out cosine value for a given cell type; negative values indicate that eNE shifted the transcriptome in the direction opposite to aging. Values from the individual held-out iterations are shown in **Extended Data Fig. 10b**.

##### Bias-corrected cosine design

As an independent estimate of buffering that uses all mice, a bias-corrected cosine was computed from group means of size-factor-normalized log2(counts + 1) pseudobulk expression. Here B = aged HC − young HC and E = aged eNE − aged HC at each age, computed over genes with ≥ 10 counts in ≥ 2 samples; cell types with fewer than 1,000 such genes were not analyzed. Noise in the shared aged-HC mean enters B and E with opposite signs and pushes the cosine negative. Its expected contribution, the variance among aged HC mice divided by their number (s²ᵍ/n), was therefore added back to the dot product for each gene, leaving the denominator uncorrected.

A null distribution was generated from 50 simulations with no aging or eNE effect, in which every mouse was drawn from a normal distribution with the gene’s overall mean and pooled within-group standard deviation. A cell type was classified as buffering at a given age if its held-out cosine was < 0 and its bias-corrected cosine fell below the null mean − 2 s.d. Middle and old age were classified separately. Buffered and amplified genes were called only in cell types that were classified as buffering at that same age. Transcription factor activity was analyzed at old age in every cell type classified as buffering at middle age, at old age, or at both, so that a cell type that buffered only at middle age still contributed its old-age transcription factor results (**Fig. 6**).

##### Buffering Index

To summarize how far eNE moved aged animals toward the young state, a buffering index (BI) was computed for each sex and cell type from the full-group DESeq2 contrasts, across genes with log_2_FC estimates in both:

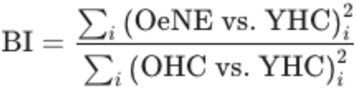

OeNE vs. YHC and OHC vs. YHC were used at old age, and MeNE vs. YHC and MHC vs. YHC at middle age. BI < 1 indicates that eNE mice were closer to young HC mice than were age-matched HC mice. Noise in the shared young HC reference enters the numerator and denominator alike and biases the BI toward 1, which is conservative, so the BI was reported without bias correction. The BI was used descriptively, plotted against the held-out cosine (**Fig. 5d**) and shown per cell type in **Extended Data Fig. 10e**, and not as a classification criterion.

##### Buffered and amplified genes

Within each cell type, genes were classified from the three held-out iterations using nominal DESeq2 Wald *p* values; because the aim was candidate discovery, no multiple-testing correction was applied at this step. In each iteration, a gene was considered aging-associated if its aging contrast had *p* < 0.05, and opposed if it was aging-associated and its eNE log_2_FC had the opposite sign to its aging log2FC. Buffered genes were defined at two stringencies: Tier 2, opposed in at least two of the three iterations, and Tier 1 (strict), opposed in all three iterations with an eNE change of *p* < 0.05 in at least two. Amplified genes were defined at the same two tiers, but with the eNE change in the same direction as the aging change. Middle and old age were classified separately. Tier 1 calls require agreement across all three aged HC mice, including the HC mouse sequenced in the same run as the eNE mice. Fold-change strips show the aging (aged HC vs. young HC) and eNE (eNE vs. HC) log_2_FC of each gene, split by the direction of the aging change. **Fig. 5c** shows all Tier 2 buffered genes, and **Extended Data Fig. 10d** names Tier 1 buffered genes; gene sets with fewer than five Tier 1 genes were not shown in **Extended Data Fig. 10d**. Pathway analyses used all Tier 2 genes.

#### Transcription factor (TF) activity inference

Upstream regulators of failed induction, basal alteration and age-emergent genes were identified by testing each gene class for over-representation of mouse CollecTRI regulons (**Fig. 3d; Extended Data Figs. 7e and 9e)**. Tests were run separately for each cell type, sex and age, and for up- and downregulated genes. For each transcription factor (TF), a one-sided Fisher’s exact test compared the fraction of its targets in the class with the fraction among all genes tested in that cell type. The background was the young eNE vs. young HC contrast for failed induction and basal alteration genes, and the aged eNE vs. aged HC contrast for age-emergent genes. Only gene sets of ≥ 5 genes and TFs with ≥ 3 targets among the tested genes were included. For each TF, enrichment was the number of its targets in the gene class divided by the number expected if targets were distributed at random, expressed as a log2 ratio. p values were adjusted with the Benjamini–Hochberg method across TFs within each test, and TFs with FDR < 0.05 were considered significant. Fisher’s test does not use the direction of regulation, so it identifies TFs whose targets are over-represented in a class, not whether the TF is activated or repressed.

#### Transcription factor regulators of buffering

TF buffering was analyzed in old mice of each sex, in cell types that passed both the held-out and bias-corrected cosine thresholds (**Fig. 6a**). TF activities were inferred with decoupleR using the univariate linear model (ULM) and signed mouse CollecTRI regulons. For each held-out iteration *k*, activities were estimated from the genome-wide Wald statistics of the same two contrasts used to define **B**ₖ and **E**ₖ, without gene thresholding. The aging activity, *a*ᴮₖ, came from the aging contrast (old HC with mouse *k* held out vs. young HC), and the eNE activity, *a*ᴱₖ, from the eNE contrast (old eNE vs. held-out old HC mouse *k*). A positive activity indicates that a TF’s activated targets increased and its repressed targets decreased relative to other genes. Activities therefore reflect coordinated changes in target expression rather than TF abundance. Each TF was assigned a buffering score, S = meanₖ(−*a*ᴮₖ × *a*ᴱₖ), averaged over the iterations that succeeded (≥ 2 of 3). S > 0 indicates that eNE shifted the TF’s activity opposite to its aging change (buffered), and S < 0 that eNE shifted it further in the aging direction (amplified).

Significance was assessed by relabeling mice. The six old mice (three HC and three eNE) can be split into two groups of three in 20 ways, and the full held-out analysis was repeated for each of the 19 alternative labelings. A TF was called buffered or amplified in a cell type if its observed score ranked within the top 5 of its 20 labelings (empirical p ≤ 0.25). With n = 3 mice per group, these calls nominate candidate regulators rather than individually significant findings. In **Fig. 6a**, TFs were ranked by their buffering score summed across the cell types in which they were called, and the aging and eNE activities shown are from the cell type with the highest score. Each activity is annotated with the median of its ULM FDR across the three held-out iterations (Benjamini–Hochberg across TFs within each iteration and contrast; * FDR < 0.05, ** < 0.01, *** < 0.001). These annotations are descriptive only and were not used for calls or ranking.

#### Neuron chat and cell chat analysis

Intercellular communication was inferred separately for each sex and each of the six age × condition groups (young, middle-aged and old; HC and NE), using log-normalized expression from all nuclei of a group pooled across mice. Ligand–receptor signaling among all annotated cell types was inferred with CellChat (v1.6.1) using the mouse CellChatDB. Over-expressed ligands and receptors were identified per cell type, communication probabilities were computed with the truncated-mean method (computeCommunProb, type = “triMean”), interactions involving cell types with fewer than 10 nuclei were removed (filterCommunication, min.cells = 10), and probabilities were aggregated into a cell type × cell type weight matrix (aggregateNet). Neurotransmitter- and neuropeptide-mediated communication among neuronal cell types (DG, CA3, CA1, CA2 and subicular excitatory neurons, and interneurons) was inferred with NeuronChat (v1.0.0) using the mouse NeuronChat database (run_NeuronChat, 100 permutations; neuronal cell types with ≥10 nuclei), and interaction strengths were aggregated across ligand–target pairs (net_aggregation, method = “weight”). Total communication strength for each group was calculated as the sum of all entries of the aggregated weight matrix. Because each network was built from nuclei pooled across mice, these values were reported descriptively, without statistical testing. (**Extended Data Fig. 10g**).

#### Comparison of chemogenetic activation with eNE

The DREADD dataset (old and young females, C21 vs. PBS) was analyzed separately for the covariate and downsampling approaches, with neurons restricted to cells with ≥ 50 mScarlet UMIs. Buffering was calculated exactly as for eNE, with old PBS mice in place of old HC and old C21 mice in place of old eNE. The aging contrast was old PBS vs. young PBS, and the stimulation contrast was old C21 vs. the held-out old PBS mouse. Held-out cosines, bias-corrected cosines and the buffering index (old C21 vs. young PBS relative to old PBS vs. young PBS) were computed for each cell type, and cell types were classified with the same thresholds (**Extended Data Fig. 10f**).

At the TF level, DREADD buffering scores were calculated for every TF and cell type in the same way as for eNE, using DREADD’s own aging axis: −(DREADD aging activity × DREADD stimulation activity). Each TF’s DREADD buffering score was plotted against its eNE buffering score in old females, separately for neurons (DG, CA3, CA1, CA2 and subicular excitatory neurons, and interneurons) and glia. The ten TFs per cell type with the highest eNE buffering score are highlighted (**Fig. 6b, downsampling; Extended Data Fig. 11a, covariate**). TFs in the upper-right quadrant were buffered by both eNE and DREADD activation. As a complementary view, DREADD buffering scores were also calculated on the eNE aging axis, −(eNE aging activity × DREADD stimulation activity), which asks whether DREADD activation moves each TF against its eNE-defined aging change. The five TFs per cell type with the highest eNE buffering score and the five with the strongest eNE amplification are highlighted (**Extended Data Fig. 11b**).

#### Statistical analysis and software

All statistical analyses were performed in R (v4.4.0; Bioconductor v3.20). Single-cell data were processed with Seurat (v5.6.0) and BPCells (v0.3.1). Ambient RNA and doublets were removed with SoupX (v1.6.2) and scDblFinder (v1.20.2). Pseudobulk differential expression used DESeq2 (v1.46.0). Neuronal activity scores were computed with neuroestimator (v0.1). TF activity was inferred with decoupleR (v2.12.0) using mouse CollecTRI regulons (1,106 TFs, 39,521 interactions), retrieved through OmnipathR (v3.14.0) on 31 July 2026. Pathway enrichment used clusterProfiler (v4.14.6) with org.Mm.eg.db (v3.20.0) and GO.db (v3.20.0). Cell–cell communication was inferred with CellChat (v1.6.1) and NeuronChat (v1.0.0). Unless otherwise stated, multiple comparisons were corrected with the Benjamini–Hochberg procedure (padj < 0.05). Buffering gene calls used nominal p values replicated across held-out iterations, and TF calls used ranks against relabeled mice, as described above. Each sex was analyzed separately. Method-specific statistical approaches, including the tests for dynamic range compression, are described in the relevant sections above. Sample sizes, sequencing runs and cell numbers for each sequenced mouse are provided in **Supplemental Table 1**; sample sizes for histological experiments are given in the figure legends. Method-specific statistical approaches, including the tests for dynamic range compression, are described in the relevant sections above; statistical tests for histological quantifications are given in the corresponding figure legends.

## Use of AI

This work made use of AI for stress-testing code, code verification, and grammatical editing.

## Data availability

Raw and processed single-cell RNA-seq data will be deposited in the NCBI Gene Expression Omnibus (GEO); submission pending. All other data supporting the findings of this study are available within the paper and its Supplementary Information. All other data requests will be made available from the authors upon reasonable request.

## Code availability

All custom analysis scripts used in this study will be made available at GitHub (https://github.com/KoreanUm97/Um2026-experience-buffers-aging-transcriptome).

## Author Contributions

J.U., C.L., and E.A.P. conceptualized and designed the study. J.U. and C.L. performed the environmental enrichment experiments. J.U. and E.A.P. performed the chemogenetic experiments. J.U. and C.L. prepared the single-cell Flex libraries. J.U. performed the single-cell analyses. E.T. integrated the single-cell datasets using GTAC@MGI computational resources and advised on the computational analyses. J.U., C.L., and K.N. performed and quantified the immunohistochemistry and RNAscope experiments. J.W.A. created graphical schematics. J.U. and E.A.P. wrote the manuscript.

## Acknowledgements

We thank Joey and the WashU OFMD fabrication team for building the enhanced novel environment cages and Jennifer Ponce and the Genome Technology Access Center at the McDonnell Genome Institute (GTAC@MGI) for sequencing. We thank members of the Pollina laboratory and J.U.’s thesis committee for critical feedback on project design, and Ava Carter, Eric Griffith, Andrew Yoo, Shin Imai, Brian Lananna, and Ian Vogel for feedback on the manuscript.

## Funder Information Declared

We acknowledge NIH grants R00AG064042 and R01AG098482 to E.A.P and the Dean’s Graduate Fellowship to J.U.

