## Extended Data Figures 1-11 for "Aging remodels the neuronal activity-dependent transcriptome"

Extended Data Figure 1

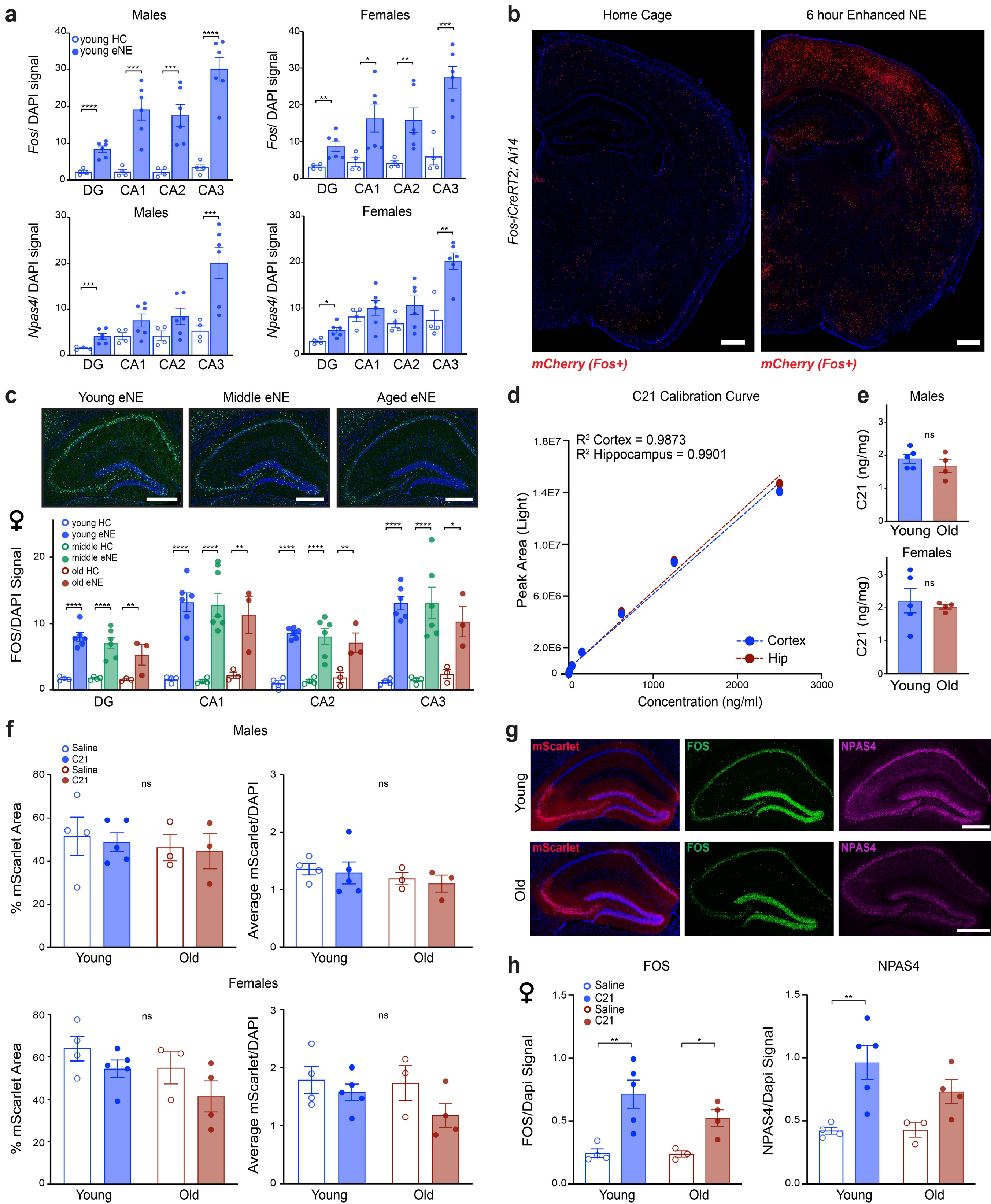

**<u>Extended Data Figure 1: Validation of activity paradigms to assess activity-dependent gene regulation across aging</u>**

a. Mean intensity ± SEM *Fos* and *Npas4 in situ* RNA signal in young males and females following 6 hour exposure to eNE. *p<0.05, **p<0.01,*** p<0.005, **** p<0.0001; p-values by 2-way ANOVA (age x stimulation) with Tukey's multiple hypothesis correction performed within each subregion (*i.e.* DG, CA1, CA2, CA3).

b. Representative images of *Fos-iCreERT2; Ai14* exposed to either home cage or 6 hours of eNE and treated with 4-OHT at the onset of exposure to eNE. mCherry+ cells turned on FOS during the 6 hour eNE exposure. Scale bar indicates 500 µm.

c. Upper: Immunostaining for FOS protein for young (4-month), middle age (14-month), or old (24 month) female mice following 6 hours of exposure to enhanced NE. Lower: Mean intensity ± SEM of FOS signal. Each dot represents a single animal as a biological replicate (n=3-6 replicates per group). *p<0.05, **p<0.01, *** <0.005, **** p<0.0001; p-values by 2-way ANOVA (age x stimulation) with Tukey's multiple hypothesis correction performed within each subregion. Scale bar indicates 500 µm.

d. Peak intensity for C21 by mass spectrometry across a dosage curve in cortex vs. hippocampus of the same mice. Cortical C21 and hippocampal abundance is closely matched and dose responsive.

e. Quantifications of C21 in cortex of young vs. old male and female mice, normalized to internal standard. p-value NS by two-sided, Welch's t-test. See **Extended Data Fig. 8b** for samples specifically used in the single cell RNA-seq.

f. Upper panel left: Mean ± SEM percentage of mScarlet (representing DREADD viral infection) per hippocampus in young vs. old male mice. Each point represents an individual animal. Upper right panel: Mean ± SEM mScarlet intensity per hippocampus in young vs. old male mice. Lower panel left: Mean ± SEM percentage of mScarlet per hippocampus in young vs. old female mice. Lower right panel: Mean ± SEM mScarlet intensity per hippocampus in young vs. old female mice. p-values NS by 2-way ANOVA (age × stimulation) with Tukey's multiple hypothesis testing.

g. Representative immunostaining of ARGs NPAS4 and FOS in young vs. old female mice injected with AAV9-hSyn-hM3D(Gq)-T2A-mScarlet-KASH. Animals were collected 2 hours post IP injection of DREADD agonist C21. Scale bar indicates 500 µm.

h. Mean intensity ± SEM of FOS (left) or NPAS4 (right) immunostaining signal in female mice. Each dot represents a single animal (n=3-5 replicates per condition). *p<0.05, ** p<0.01, *** p<0.005; p-values by 2-way ANOVA (age × stimulation) with Tukey's multiple hypothesis testing.

Extended Data Figure 2

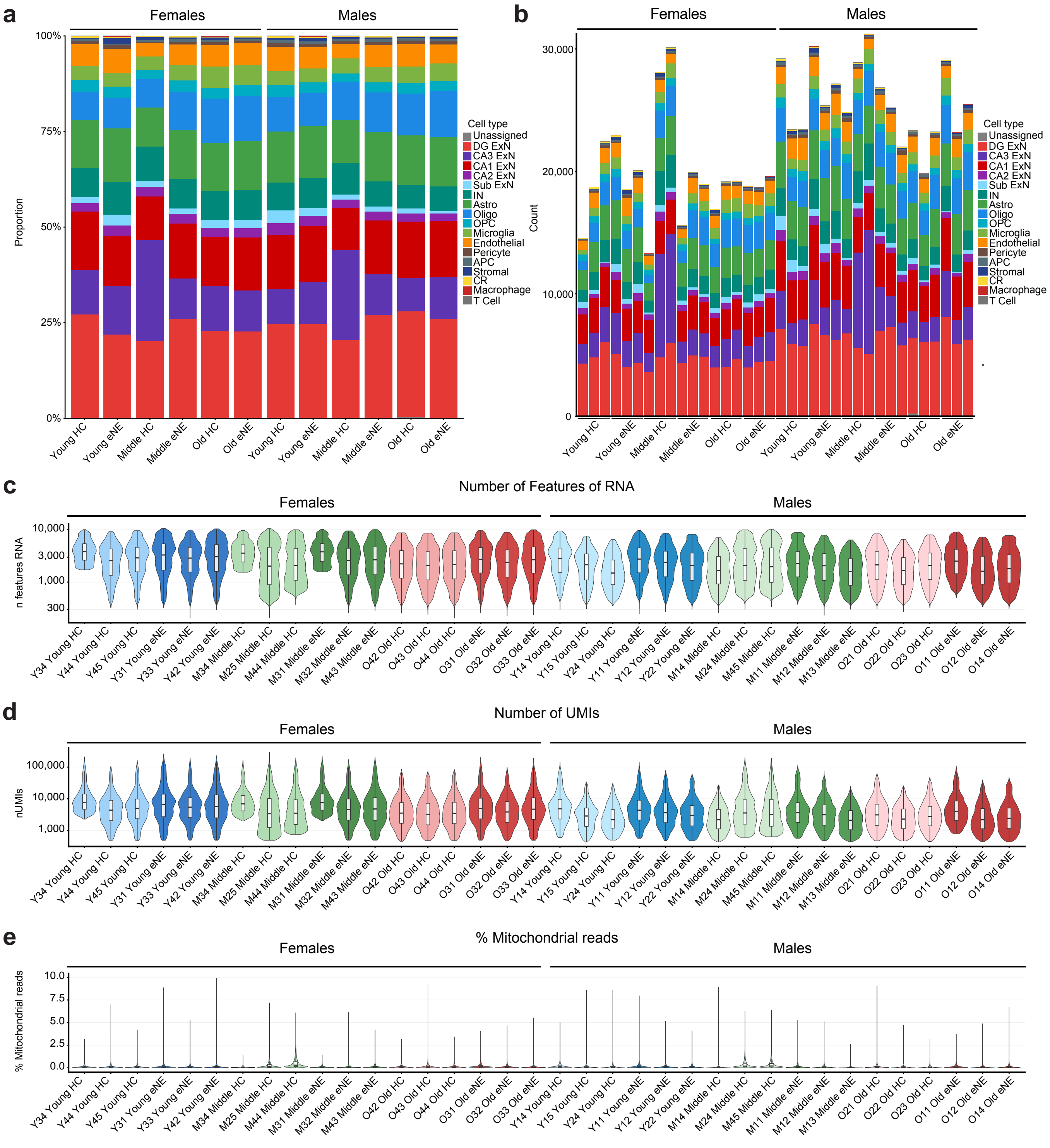

**Extended Data Figure 2: Quality control of the eNE single-cell RNA-seq dataset**

a. Proportion of each major cell type within each group (age × stimulation) for females (left) and males (right), with cells pooled across the three mice per group. ExN, excitatory neuron; IN, inhibitory neuron; Sub, subiculum; Astro, astrocyte; oligo, oligodendrocyte; OPC, oligodendrocyte precursor cell; Endo, endothelial cell; APC, astrocyte progenitor cell; CR, Cajal–Retzius cell.

b. Number of cells per cell type for each individual sample (n = 36 mice; 3 per sex × age × stimulation state), ordered by sex, age, and condition.

c-e. Per-sample distributions of the number of detected genes (**c**), number of UMIs (**d**) and percentage of mitochondrial reads (**e**). Violins show the distribution across cells, and inner boxplots show the median and interquartile range. Samples are colored by age (blue, young; green, middle-aged; red, old) and condition (light, HC; dark, eNE). The y-axes in **c** and **d** are $\log_{10}$-scaled.

Extended Data Figure 3

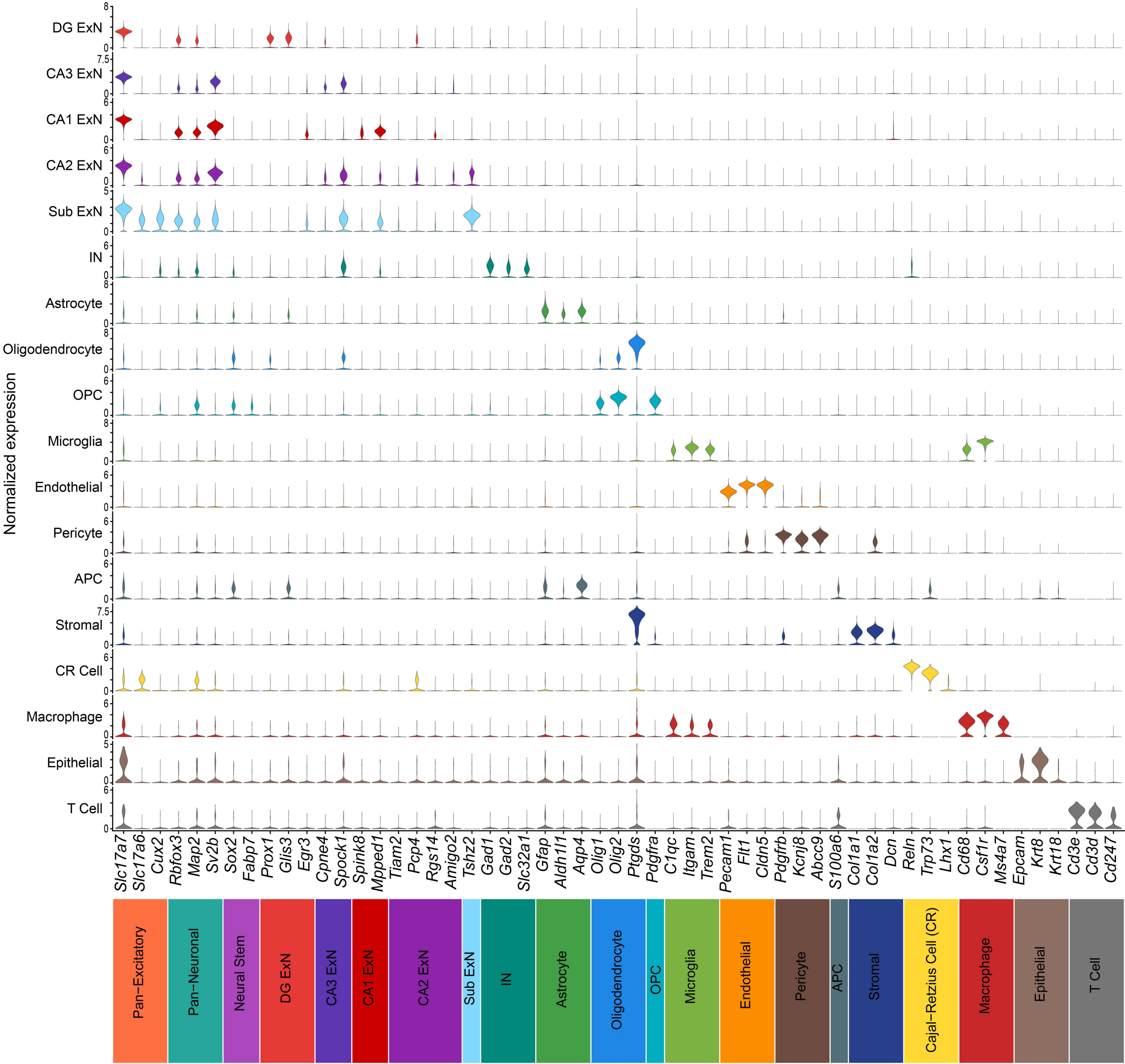

**Extended Data Figure 3: Canonical marker gene expression across hippocampal cell types**

a. Stacked violin plots of log-normalized expression of canonical marker genes (columns) across the 18 hippocampal cell types (rows) identified in the integrated eNE and HC dataset (n = 36 mice). Markers are grouped by the population they identify: pan-neuronal (*Rbfox3*, *Map2*, *Sv2b*); pan-excitatory (*Slc17a7*, *Slc17a6*, *Cux2*); neural stem and progenitor (*Sox2*, *Fabp7*); dentate gyrus excitatory neurons (*Prox1*, *Glis3*, *Egr3*); CA3 excitatory neurons (*Cpne4*, *Spock1*); CA1 excitatory neurons (*Spink8*, *Mpped1*, *Tiam2*); CA2 excitatory neurons (*Pcp4*, *Rgs14*, *Amigo2*); subiculum excitatory neurons (*Tshz2*); inhibitory neurons (*Gad1*, *Gad2*, *Slc32a1*); astrocytes (*Gfap*, *Aldh1l1*, *Aqp4*); oligodendrocytes (*Olig1*, *Olig2*, *Ptgds*); oligodendrocyte precursor cells (*Pdgfra*); microglia (*C1qc*, *Itgam*, *Trem2*); endothelial cells (*Pecam1*, *Flt1*, *Cldn5*); pericytes (*Pdgfrb*, *Kcnj8*, *Abcc9*); astrocyte progenitor cell (APC)(*S100a6*); stromal cells (*Col1a1*, *Col1a2*, *Dcn*); Cajal–Retzius (CR) cells (*Reln*, *Trp73*, *Lhx1*); macrophages (*Cd68*, *Csf1r*, *Ms4a7*); epithelial cells (*Epcam*, *Krt8*, *Krt18*); and T cells (*Cd3e*, *Cd3d*, *Cd247*). Violin width reflects the distribution of expression across cells within each cell type. See Methods for full panels of markers used.

Extended Data Figure 4

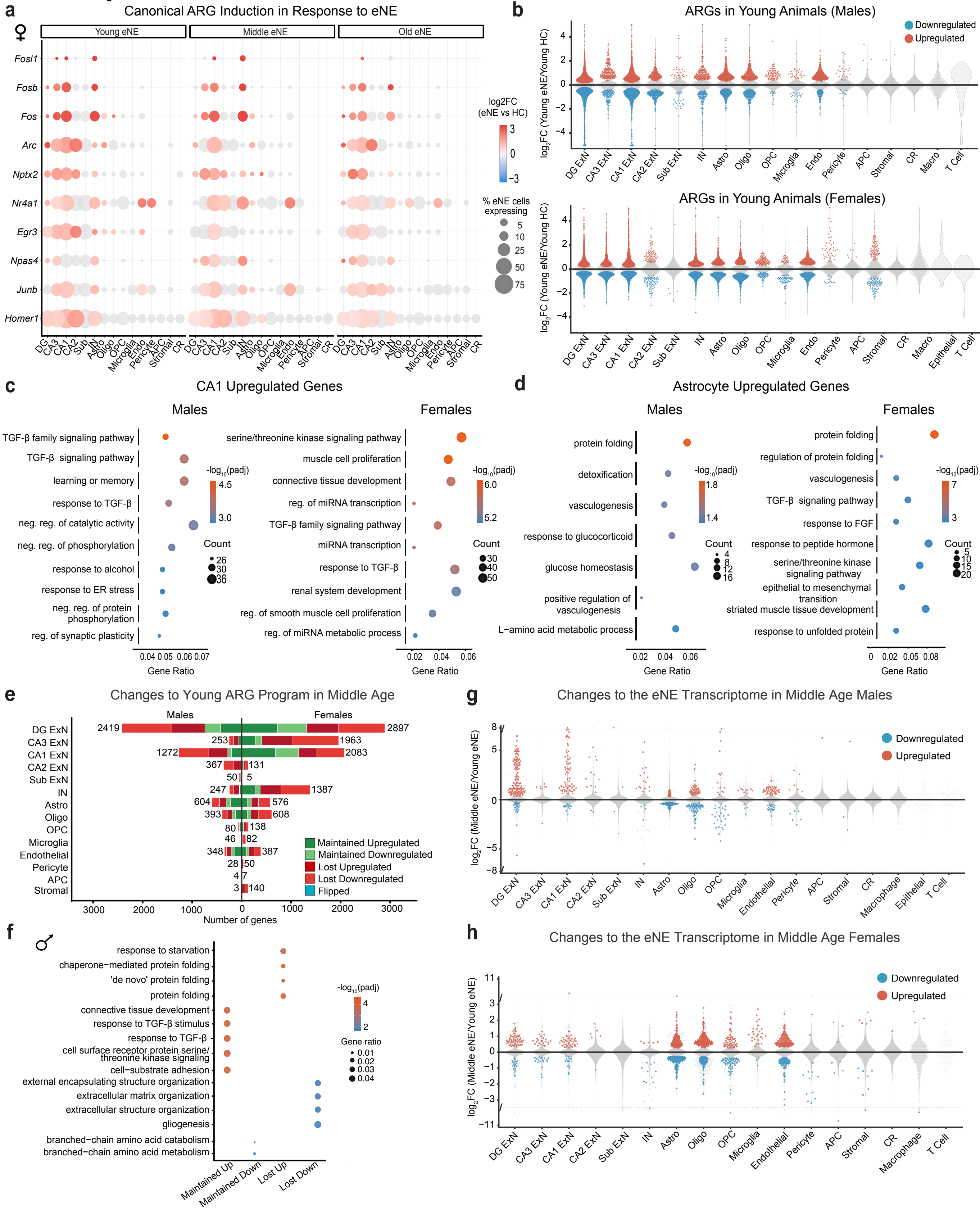

**Extended Data Figure 4: Age-dependent remodeling of the ARG transcriptome**

a. Dot plot of canonical ARGs across cell types in young, middle-aged, and old female mice. Dot color indicates $\log_2$ fold change (eNE vs. HC), and dot size indicates the percentage of eNE cells expressing the gene. Grey dots indicate non-significant changes. Males are shown in **Fig. 2b**.

b. Strip volcano plots of differential expression between young eNE vs. young HC mice per cell type in males (top) and females (bottom). Each point represents a gene. Red and blue indicate significantly upregulated and downregulated genes, respectively ($padj < 0.05$); grey indicates non-significant genes.

c,d. Gene Ontology (Biological Process) enrichment of genes upregulated by eNE in young CA1 ExNs (**c**) and young astrocytes (**d**), in males (left) and females (right). Dot size indicates gene count, color indicates $-\log_{10}(padj)$, and the x-axis shows gene ratio.

e. Fate of young ARGs in middle age per cell type (males left, females right). Young ARGs were classified by their regulation in middle eNE vs. middle HC mice: Maintained Upregulated and Maintained Downregulated (significant in the same direction in middle age), Lost Upregulated and Lost Downregulated (no longer significant, $padj \geq 0.05$), and Flipped (significant in the opposite direction in young vs. old). Numbers indicate the total number of young ARGs per cell type.

f. Gene Ontology (Biological Process) enrichment of Maintained and Lost ARG classes in old males, pooled across cell types. Dot color indicates $-\log_{10}(padj)$, and dot size indicates gene ratio. Females are shown in **Fig. 2f.**

g,h. Strip volcano plots of differential expression between middle-aged eNE vs. young eNE mice per cell type in males (**g**) and females (**h**). Genes significantly different between middle-aged HC and young HC mice were excluded. ExN, excitatory neuron; IN, inhibitory neuron; Sub, subiculum; Astro, astrocyte; oligo, oligodendrocyte; OPC, oligodendrocyte precursor cell; Endo, endothelial cell; Macro, macrophage; APC, astrocyte progenitor cell; CR, Cajal–Retzius cell.

Extended Data Figure 5

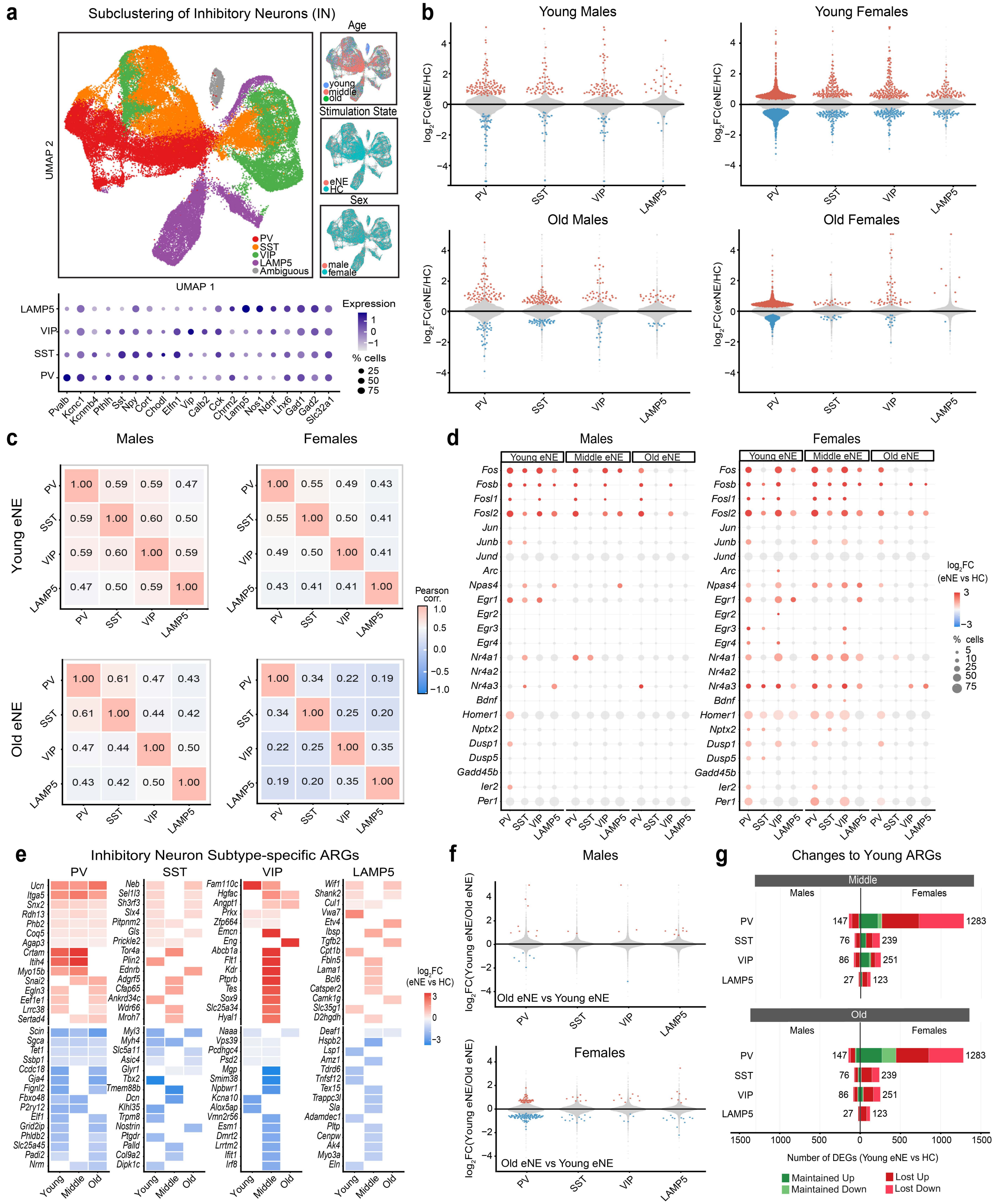

**Extended Data Figure 5: Activity-dependent transcription in inhibitory neuron subtypes**

a. UMAP of subclustered inhibitory neurons (INs) colored by subtype (PV, SST, VIP, LAMP5 and ambiguous), with insets colored by age, stimulation state and sex (top). Dot plot of subtype marker gene expression (bottom). Dot color indicates scaled average expression and dot size the percentage of cells expressing the gene. Ambiguous cells were excluded from downstream analyses.

b. Strip volcano plots of differential expression between eNE vs. HC mice per IN subtype in young (top) and old (bottom) males (left) and females (right). Each point represents a gene. Red and blue indicate significantly upregulated and downregulated genes, respectively (padj < 0.05); grey indicates non-significant genes.

c. Pearson correlation of eNE vs. HC $log_2FC$ profiles across all tested genes between IN subtypes in young (top) and old (bottom) males (left) and females (right).

d. Dot plot of canonical ARGs across IN subtypes in young, middle-aged, and old males (left) and females (right). Dot color indicates $log_2FC$ (eNE vs. HC) for significant genes (padj < 0.05); grey dots indicate non-significant genes. Dot size indicates the percentage of eNE cells expressing the gene.

e. Heat map of subtype-specific ARGs, pooled across sexes. These were genes significantly regulated by eNE (padj < 0.05) in one IN subtype but not in any other IN subtype or any excitatory neuron population at the same age. Color indicates $log_2$ fold change (eNE vs. HC) at each age. Within each subtype, upregulated genes are shown above and downregulated genes below. Shown are the top 15 genes per direction, ranked by the number of ages at which the gene was subtype-specific and then by maximum $| log_2FC |$.

f. Strip volcano plots of differential expression between old eNE vs. young eNE mice per IN subtype in males (top) and females (bottom). Colors are as in **b**.

g. Fate of the young ARG program at middle age (top) and old age (bottom) per IN subtype (males left, females right). Young ARGs were classified as Maintained Up or Maintained Down (significant in the same direction in older animals) or as Lost Up or Lost Down (no longer significant). Because the gene set is anchored in young mice, the total per subtype is the same at both ages; only the proportion maintained and lost differs.

Extended Data Figure 6

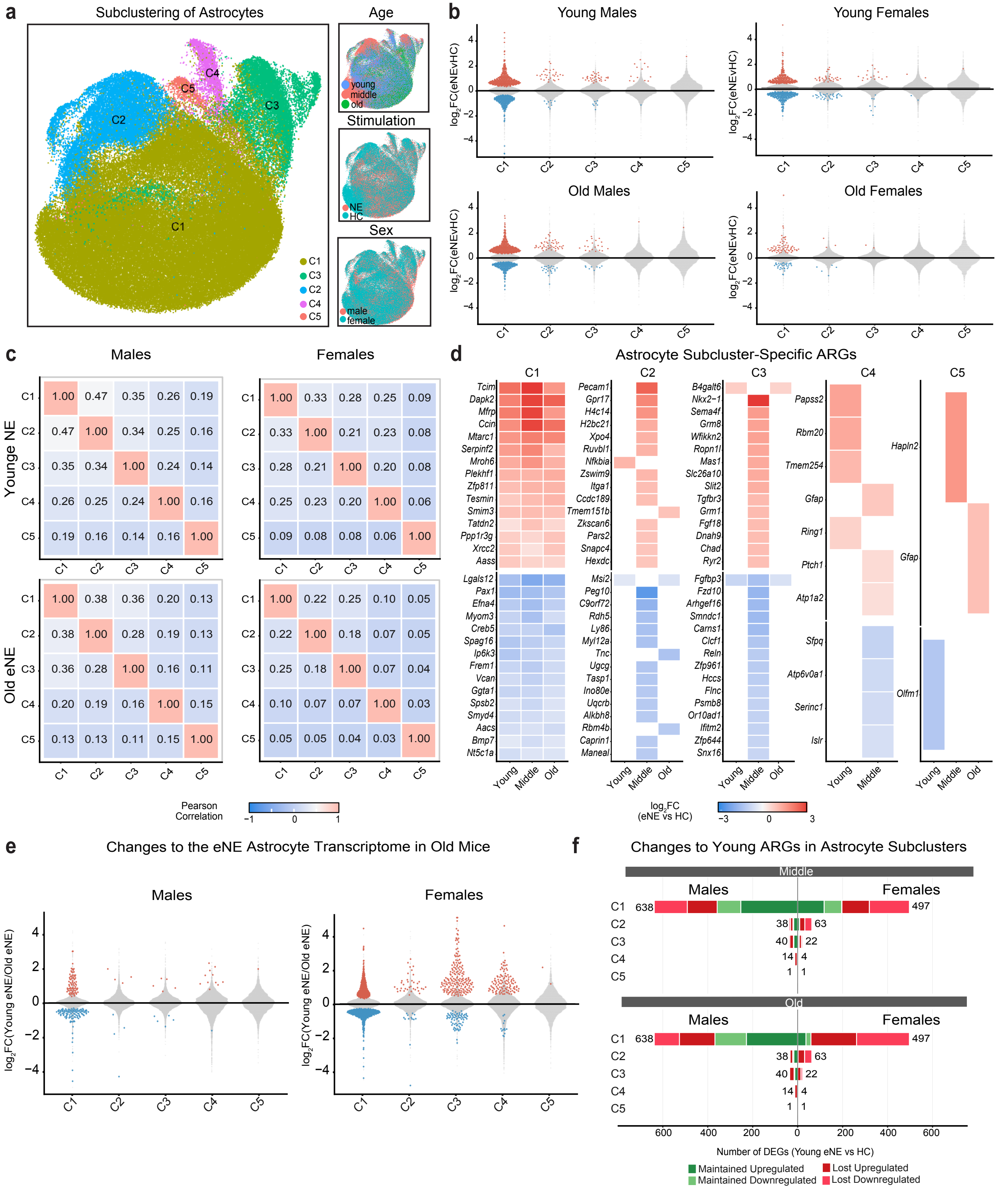

**Extended Data Figure 6: Activity-dependent transcription in hippocampal astrocyte subclusters**

a. UMAP of subclustered astrocytes colored by subcluster (C1–C5), with insets colored by age, stimulation state, and sex.

b. Strip volcano plots of differential expression between eNE and HC mice per astrocyte subcluster in young (top) and old (bottom) males (left) and females (right). Each point represents a gene. Red and blue indicate significantly upregulated and downregulated genes, respectively ($padj < 0.05$); grey indicates non-significant genes.

c. Pearson correlation of eNE vs. HC $log_2FC$ profiles across all tested genes between astrocyte subclusters in young (top) and old (bottom) males (left) and females (right).

d. Heat map of subcluster-specific ARGs, pooled across sexes. Shown are genes regulated by eNE ($padj < 0.05$) in one astrocyte subcluster but not in any other subcluster, any excitatory neuron population, or inhibitory neuron at the same age. Color indicates $log_2$ fold change (eNE vs. HC) at each age. Shown are the top 15 genes per direction, ranked by the number of ages at which the gene was subtype-specific and then by maximum $| log_2FC |$.

e. Strip volcano plots of differential expression between old eNE and young eNE mice per astrocyte subcluster in males (left) and females (right). Colors are as in **b**.

f. Fate of the young ARG program at middle age (top) and old age (bottom) per astrocyte subcluster (males left, females right). Young ARGs were classified as Maintained Up or Maintained Down (significant in the same direction at the older age) or as Lost Up or Lost Down (no longer significant, $padj \geq 0.05$). Because the gene set is anchored in young mice, the total per subcluster is the same at both ages; only the proportion maintained and lost differs.

Extended Data Figure 7

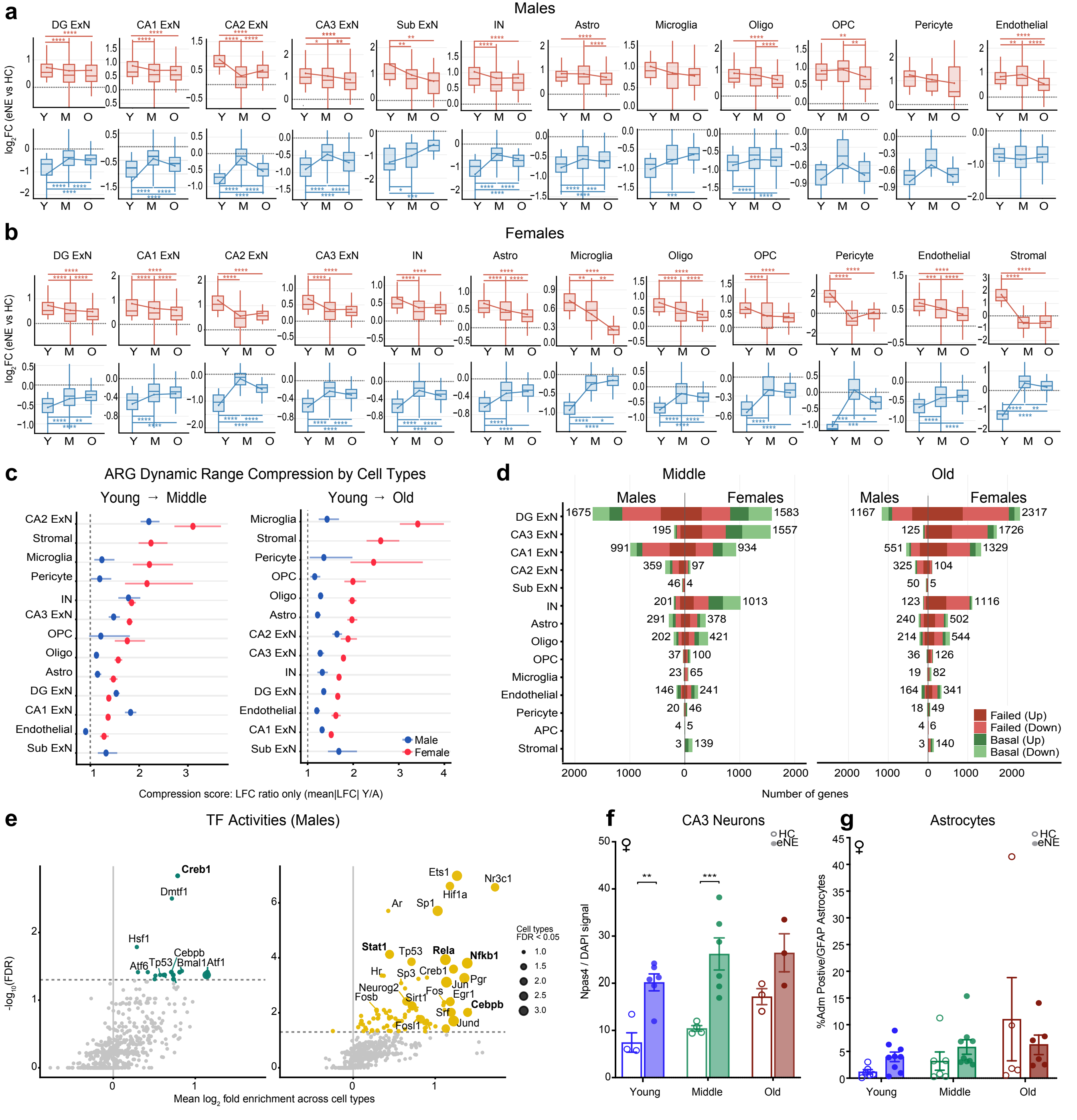

**Extended Data Figure 7: Dynamic range compression and modes of ARG failure across hippocampal cell types and sexes**

a,b. Per-gene $log_2FC$ (eNE vs. HC) of young ARGs in young (Y), middle-aged (M), and old (O) males (**a**) and females (**b**) across cell types. Young ARGs were genes significantly upregulated (top rows) or downregulated (bottom rows) in young eNE relative to young HC mice (padj < 0.05), and their $log_2FC$ was tracked at middle and old age. Boxplots show the median and interquartile range, with whiskers extending to 1.5× IQR and outliers not displayed. The y-axis is truncated at the 95th percentile for display; statistics were computed on all genes. **p* < 0.05, ***p* < 0.01, ****p* < 0.001, *****p* < 0.0001; two-sided, paired Wilcoxon signed-rank test on all genes in the set, Benjamini–Hochberg corrected for multiple age comparisons within each cell type. Only cell types with ≥ 10 young ARGs in a given direction are shown.

c. Dynamic range compression score per cell type for the young-to-middle (left) and young-to-old (right) transitions, with males (blue) and females (red). For each cell type, the compression score for Young ARGs was calculated as the mean $|log_2FC|$ in young divided by the mean |log2FC| of those same genes at the older ages. Scores were averaged across the upregulated and downregulated sets. Horizontal lines show 95% confidence intervals from 1,000 bootstrap resampling of genes. The dashed line marks a score of 1 (no compression); values above 1 indicate a weaker eNE response with age. Cell types are ordered by the mean score across sexes.

d. Classification of lost young ARGs at middle age (left) and old age (right) per cell type (males left, females right). Amongst these lost genes, basal-alteration genes were changed at rest (middle or old HC vs. young HC) by at least half the young eNE magnitude. All other lost genes were classified as failed induction (Up) or failed repression (Down). Numbers indicate the total number of lost genes per cell type.

e. CollecTRI transcription factor regulons enriched among failed-induction (left) and basal-alteration (right) genes in males, plotted as in **Fig. 3d**. Dot size indicates the number of cell types with FDR < 0.05, and the dashed line marks that threshold.

f,g. *Npas4* signal normalized to DAPI in CA3 neurons (**f**) and percentage of $GFAP^+$ astrocytes positive for *Adm* (**g**) in young, middle-aged, and old female HC (open) and eNE (filled) mice. Data are mean ± SEM, and each dot represents one mouse. ***p* < 0.01, ****p* < 0.001; two-way ANOVA (age × stimulation) with Tukey's multiple comparisons test.

Extended Data Figure 8

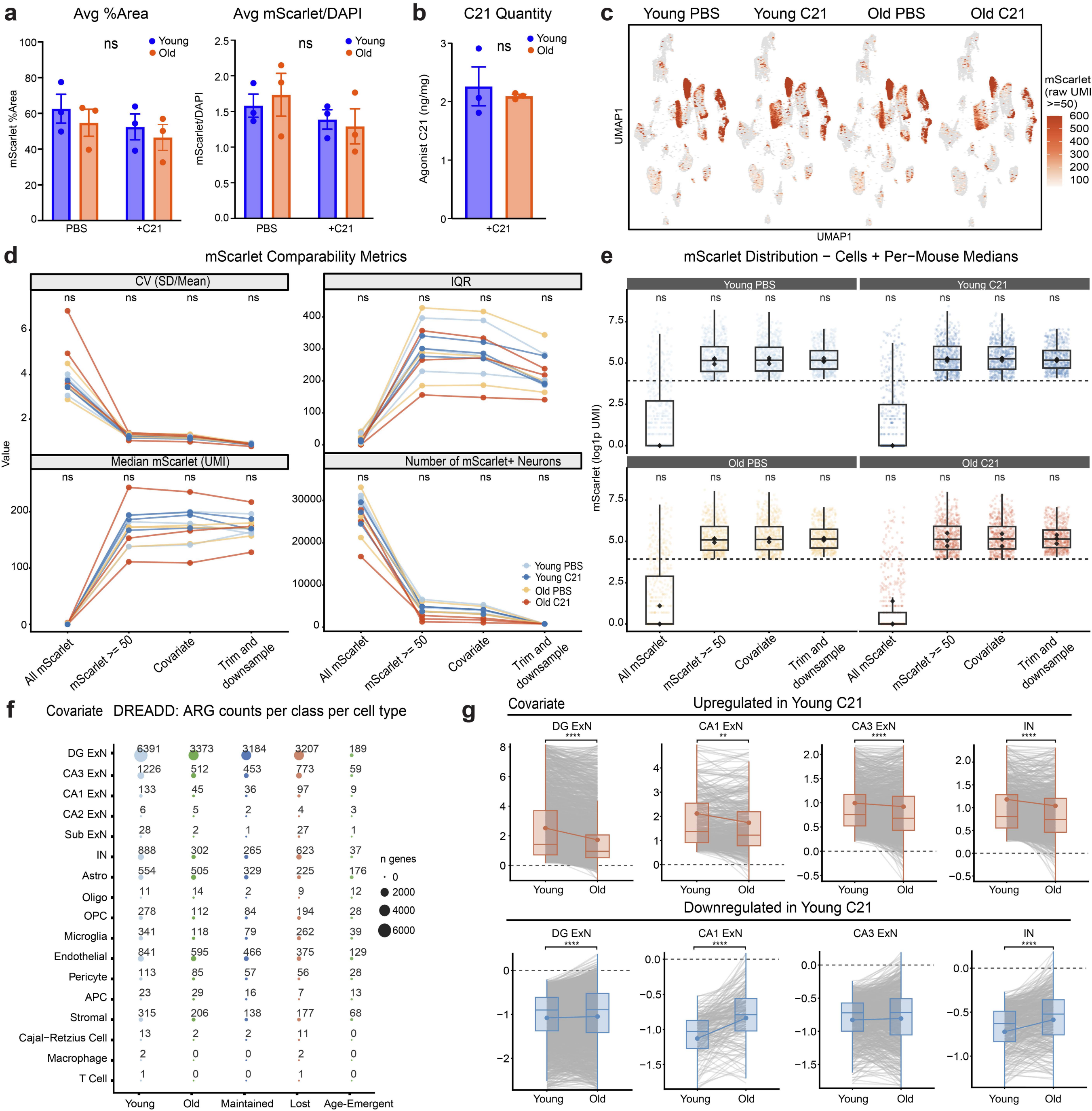

**Extended Data Figure 8: Validation of the DREADD single-cell dataset and analysis with mScarlet covariate correction**

a. Mean mScarlet-positive area (left) and mean mScarlet intensity normalized to DAPI (right) per hippocampus in young and old female mice treated with C21 or PBS (mean ± SEM). Data is re-plotted from **Extended Data Fig. 1** for the animals specifically used in single-cell RNA-seq. Each dot represents a single animal. p-value NS by two-way ANOVA (age × stimulation) with Tukey's multiple comparisons test.

b. C21 concentration in cortex of young and old C21-treated mice, measured by mass spectrometry (mean ± SEM). Data re-plotted from **Extended Data Fig. 1** for the animals used in single-cell RNA-seq. Each dot represents a single animal. NS; two-sided Welch's t-test.

c. UMAP of DREADD samples split by group, colored by mScarlet expression in cells with ≥ 50 raw mScarlet UMI.

d. Per-mouse mScarlet comparability metrics in neurons across the four groups at each processing step: all cells, the ≥ 50 UMI threshold, covariate correction, and trimming plus downsampling. Metrics shown are coefficient of variation, interquartile range, median mScarlet UMI and number of mScarlet-positive neurons. Each line represents one mouse.

e. Distribution of neuronal mScarlet expression ($\log_1$p UMI) at each processing step, per group. Points represent cells and diamonds represent per-mouse medians. The dashed line marks the 50 UMI threshold. In **d** and **e**, NS indicates no significant difference across groups by Kruskal–Wallis test on per-mouse values.

f. Number of ARGs per cell type with covariate correction, in five classes: genes significantly regulated by C21 in young mice (Young) or old mice (Old); genes significant at both ages (Maintained); genes significant in young but not old mice (Lost); and genes significant in old but not young mice (Age-emergent). Dot size indicates gene number.

g. Per-gene $\log_2$FC (C21 vs. PBS) of young-upregulated (top) and young-downregulated (bottom) ARGs in female DG ExNs, CA1 ExNs, CA3 ExNs and INs. Boxplots show the median and interquartile range, with whiskers extending to 1.5× IQR and outliers not displayed. Grey lines connect each gene's $\log_2$FC across ages; the y-axis is set to the boxplot range, so only genes falling within that range are drawn. $^{**}p < 0.01$, $^{***}p < 0.001$, $^{****}p < 0.0001$; two-sided, paired Wilcoxon signed-rank test on all genes in the set, Benjamini–Hochberg corrected for multiple age comparisons within each cell type.

Extended Data Figure 9

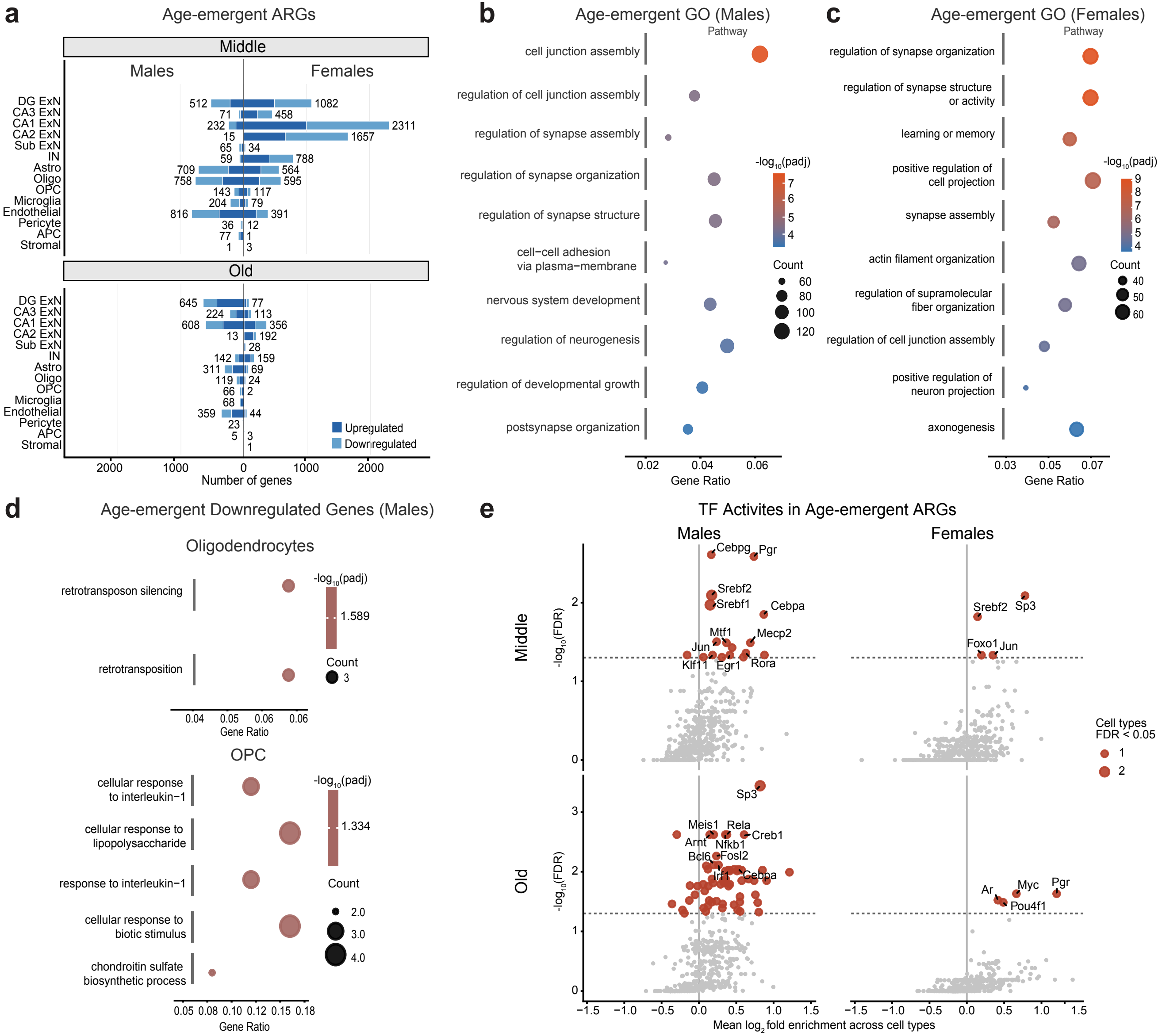

**Extended Data Figure 9: Latent transcriptional responses to activity in the aging hippocampus**

a. Number of age-emergent ARGs per cell type at middle (top) and old (bottom) age (males left, females right). Age-emergent ARGs were significantly regulated by eNE at the older age (BH-adjusted $p < 0.05$) but not in young eNE vs. HC mice. Dark and light blue indicate upregulated and downregulated genes, and numbers give the total number of age emergent genes.

b,c. Gene Ontology (Biological Process) enrichment of age-emergent ARGs in old males (**b**) and old females (**c**), pooled across cell types. Input genes included the union of up and downregulated genes. The top 10 terms by adjusted p-value are shown after merging redundant terms (semantic similarity 0.7). Color indicates $-\log_{10}(\text{padj})$, dot size indicates gene count, and the x-axis shows gene ratio.

d. Gene Ontology enrichment of age-emergent ARGs downregulated by eNE in old male oligodendrocytes (top) and OPCs (bottom), as in **b**.

e. CollecTRI transcription factor regulons enriched among age-emergent genes in males (left) and females (right), at middle (top) and old (bottom) age, plotted as in **Fig. 3d**. Dot size indicates the number of cell types with FDR < 0.05, and the dashed line marks that threshold.

Extended Data Figure 10

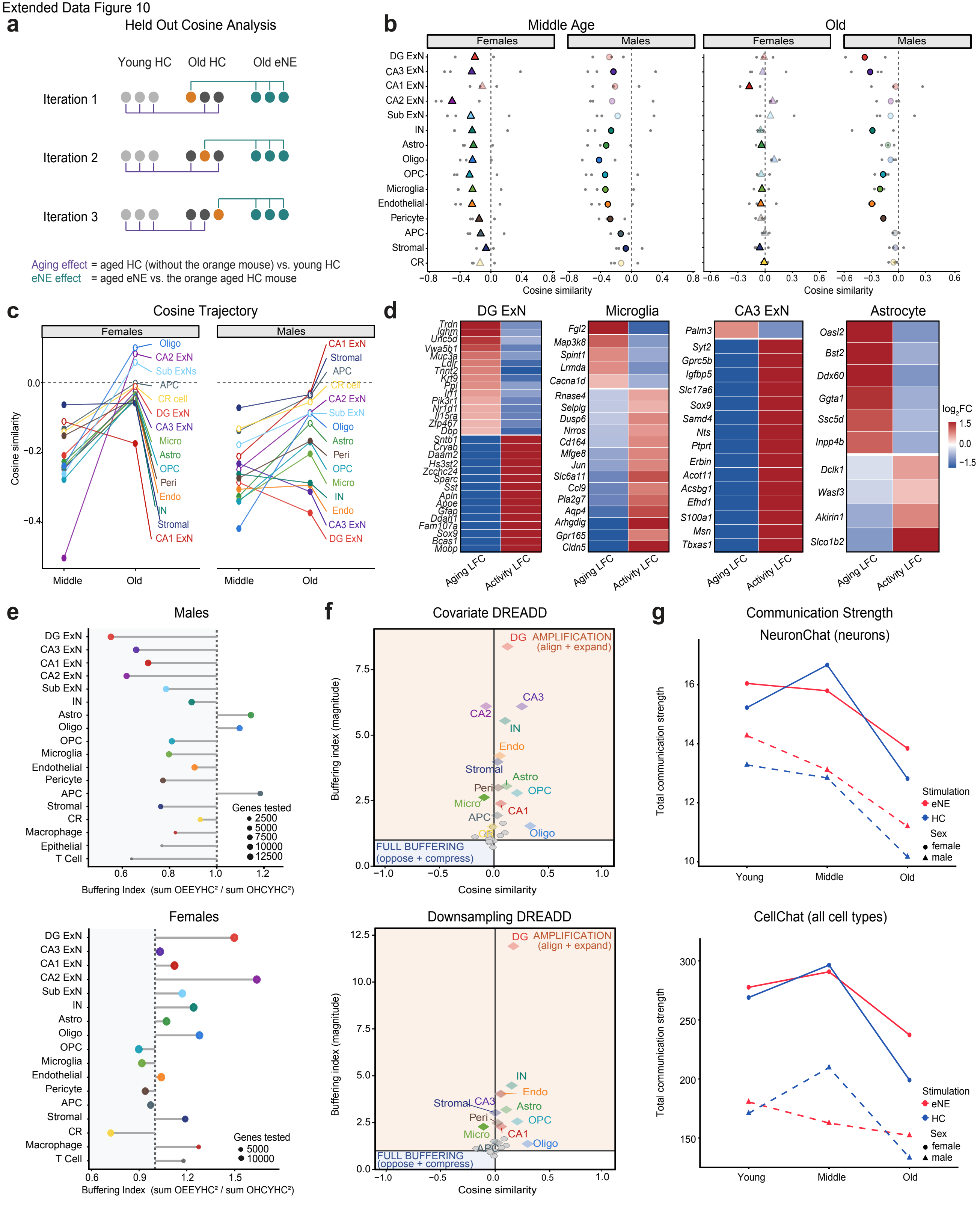

**Extended Data Figure 10: Activity-induced buffering across aged hippocampal cell types**

a. Schematic of the held-out design. In each of three iterations, one aged HC mouse (orange) was held out. The aging effect compared the remaining two aged HC mice with young HC mice, and the eNE effect compared aged eNE mice with the held-out aged HC mouse, so that no mouse contributed to both contrasts.

b. Held-out cosine similarity between the aging and eNE vectors per cell type at middle (left) and old (right) age in females (triangles) and males (circles). Small grey points are the three individual iterations, and large symbols indicate their mean. Dark symbols mark cell types that were confirmed by two independent methods (held-out cosine and bias-corrected cosine); faded symbols mark those cell types supported by only one.

c. Change in held-out cosine from middle to old age per cell type in females (left) and males (right). Filled points were confirmed by two independent methods; open points mark those cell types supported by only one.

d. Strictly buffered genes (tier T1, see **Methods**) in male DG ExNs, microglia and CA3 ExNs and female astrocytes, corresponding to **Fig. 5c**. For each gene, the left column shows the aging change (old HC vs. young HC) and the right column the eNE response in old mice (old eNE vs. old HC), as mean $\log_2$FC across held-out iterations.

e. Buffering index per cell type at old age in males (top) and females (bottom). The dashed line marks BI = 1, and BI < 1 indicates that old eNE mice are closer to young HC mice than old HC mice are. Dot size indicates the number of genes tested.

f. Held-out cosine similarity versus buffering index for DREADD activation in old females, with covariate correction (top) and trimming and downsampling (bottom). The aging vector was old PBS vs young PBS, and the stimulation vector was old C21 vs held-out old PBS mouse. $BI = \Sigma \log2FC^2$ (old C21 vs. young PBS) / $\Sigma \log2FC^2$ (old PBS vs. young PBS). Dark symbols mark cell types that were confirmed by two independent methods (held-out cosine and bias-corrected cosine); faded symbols mark those cell types supported by only one. Quadrants are as in **Fig. 5d**.

g. Total communication strength in young, middle-aged, and old mice inferred by NeuronChat for neurons (top) and by CellChat for across all cell types (bottom). HC (blue) and NE (red) in females (solid lines, circles) and males (dashed lines, triangles). Total strength was summed across all cell-type pairs for each sex × age × stimulation group.

Extended Data Figure 11

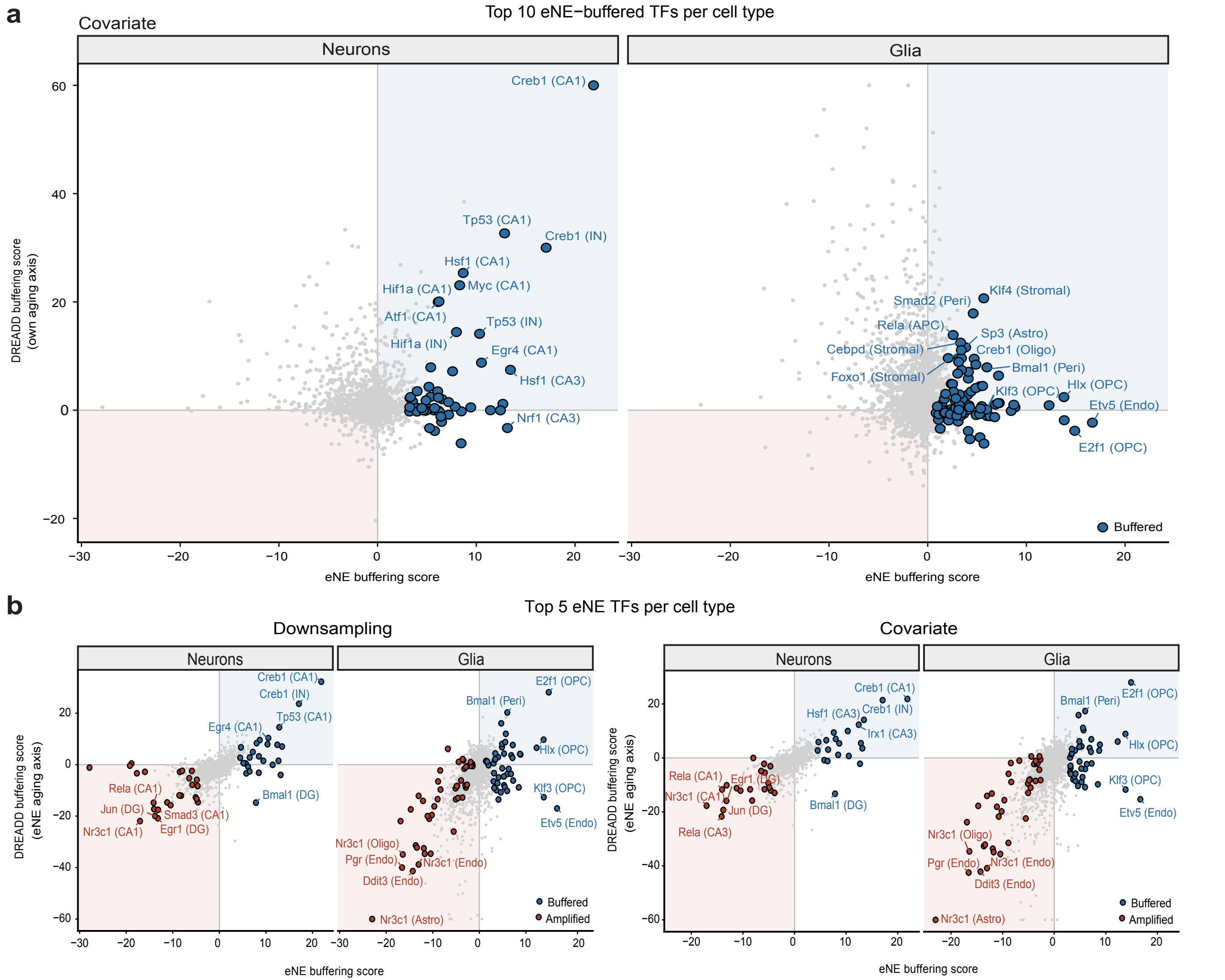

**Extended Data Figure 11: Comparison of buffering TF activity in DREADD and eNE paradigms**

a. DREADD buffering score plotted against eNE buffering score for all overlapping TFs, using the covariate approach. The ten most strongly eNE-buffered TFs per cell type are blue; all others are grey, and the strongest are labelled with their cell type. TFs in the upper-right quadrant (shaded blue) are buffered by both eNE and DREADD; those in the lower-left (shaded red) are amplified by both.

b. As in **a**, with the DREADD buffering score calculated against the eNE aging change. Here, the basal aging effect was calculated from a fully separate data set collected in the eNE dataset, for the downsampling (left) and covariate (right) approaches. This approach allows generalization across aging datasets collected independently. Highlighted are the five most eNE-buffered (blue) and five most eNE-amplified (red) TFs per cell type.
